# Fingerprinting disease-derived protein aggregates reveals unique signature of Motor Neuron Disease

**DOI:** 10.1101/2025.03.04.641150

**Authors:** Dezerae Cox, Melanie Burke, Sara Milani, Matthew A White, Fergal M Waldron, Dorothea Böken, Evgeniia Lobanova, Jemeen Sreedharan, Jenna M Gregory, David Klenerman

**Affiliations:** Yusuf Hamied Department of Chemistry, University of Cambridge, Cambridge, UK; UK Dementia Research Institute, University of Cambridge, Cambridge, UK; Molecular Horizons and School of Science, University of Wollongong, Wollongong, Australia; Maurice Wohl Clinical Neuroscience Institute, Department of Basic and Clinical Neuroscience, Institute of Psychiatry, Psychology and Neuroscience, King’s College London, London, UK; Institute of Medical Sciences, University of Aberdeen, Aberdeen, UK

## Abstract

Inappropriate aggregation of TAR DNA-binding protein 43 (TDP-43) is a hallmark of motor neuron disease (MND). Current methods for quantifying heterogeneous aggregate populations in biofluids are limited, precluding their routine use in diagnosis and disease monitoring. Single molecule microscopy methods overcome these limitations to deliver quantitative morphological and compositional fingerprinting of molecular assemblies containing TDP-43. Here, we demonstrate the application of such methods to extracts from donor brain tissues. We show the number and morphology of aggregates derived from frontal cortex and cerebellar samples is sufficient to distinguish MND donors from neurologically normal controls, as well as between different MND cohorts. In addition, we compare proteomic and microscopic compositional profiles, demonstrating how complementary insights delivered by each technique can enhance our understanding of disease mechanisms. These single molecule assays will inform future diagnostic and stratification technologies, improving our ability to deliver patient care in a timely and targeted manner.

## INTRODUCTION

Inappropriate protein aggregation is a central molecular signature of chronic neurodegenerative diseases; in Motor Neuron disease (MND), 97% of cases are reported to develop insoluble deposits of TAR DNA-binding protein 43 (TDP-43)^1^. TDP-43 is a highly conserved protein predominantly located in the nucleus, where it plays an essential role in RNA transcription, splicing, and nucleocytoplasmic transport. In MND-affected neurons, TDP-43 is sequestered in the cytoplasm by the accumulation of large insoluble aggregates decorated with aberrant post-translational modifications^2^. Mutations in TDP-43 are also a rare cause of MND, and some studies suggest that these variants increase the propensity of TDP-43 to aggregate^3^. However, decades of research are yet to satisfactorily resolve the role of aggregates in disease pathology and progression.

Existing methods for measuring disease-derived multimeric assemblies can be loosely categorised as those that quantify or characterise. Immunohistochemistry and immunofluorescence in fixed post-mortem tissue slices has long enabled the cellular and macroscopic distribution of TDP-43 to be quantified^4^, and is used for subtyping or staging of suspected MND cases^5^. Biochemical approaches, such as western blotting of aggregates extracted from solution via filter-trap or pelleting assays, have contributed to quantification of aggregate ‘load’ in cell and animal models of disease^6–8^. However, these methods are poorly suited to quantifying small soluble assemblies which likely precede the formation of large insoluble deposits, and which have been implicated in disease pathology^9,10^. More recently, cryo-electron microscopy has been instrumental in providing ultrastructural details of TDP-43 aggregates^11^, offering valuable insights into the morphological features of aggregates taken directly from disease tissues. This technology is amenable to elucidating smaller aggregate structures; however, it does not readily permit relative quantitation of these aggregates across various samples. Importantly, the ability of cryo-electron microscopy to both quantify and characterise rare subpopulations is severely limited owing to the need for particle averaging. Thus, novel quantitative approaches are urgently needed to simultaneously quantify and characterise the heterogeneity of multimeric TDP-43 assemblies in complex mixtures derived from human donors.

Single molecule microscopy methodologies offer the ability to simultaneously quantify and characterise nanoscopic multimeric assemblies from diverse sources^12–14^. However, the potential of these cutting-edge methodologies is yet to be realised in the context of MND diagnosis, stratification and treatment. Here, we report the application of a single-molecule pulldown (SiMPull) technology to characterising protein assemblies derived from post-mortem tissues of MND donors. Using this technology, we demonstrate that nanoscopic assemblies comprised of TDP-43 encode sufficient information to distinguish MND-derived samples from neurologically normal controls. These advances, combined with the flexibility and applicability of the SiMPull technology to a diverse range of biofluids, will serve as a valuable catalyst for novel mechanistic, diagnostic, and treatment studies targeting MND onset and progression.

## RESULTS

### Single-molecule pulldown (SiMPull) enables specific characterisation of TDP-43 aggregate assemblies

To detect multimeric assemblies of TDP-43, we adopted the single-molecule pulldown technology (SiMPull; Figure 1A)^15^. Like the logic of sandwich ELISA platforms, SiMPull uses antibodies tethered to a functionalised glass coverslip to capture the target of interest from solution, which is then detected using a second, fluorescently labelled antibody. Using the same monoclonal antibody for capture and detection of the target ensures the captured particle contains at least two copies of the same epitope, thereby excluding monomeric protein. The resultant fluorescence can then be quantified via total-internal fluorescence microscopy (TIRF), such that the number of target molecules is easily read out as the number of diffraction-limited fluorescent puncta. We have successfully used this strategy to specifically and sensitively detect tau assemblies derived from Alzheimer’s disease biofluids^12^. Applying the same logic, we selected an antibody (flTDP) that recognises the 203-209 residue region^16^ of TDP-43 capable of detecting full-length and truncated TDP-43 isoforms commonly associated with MND^17^.

**Figure 1:**
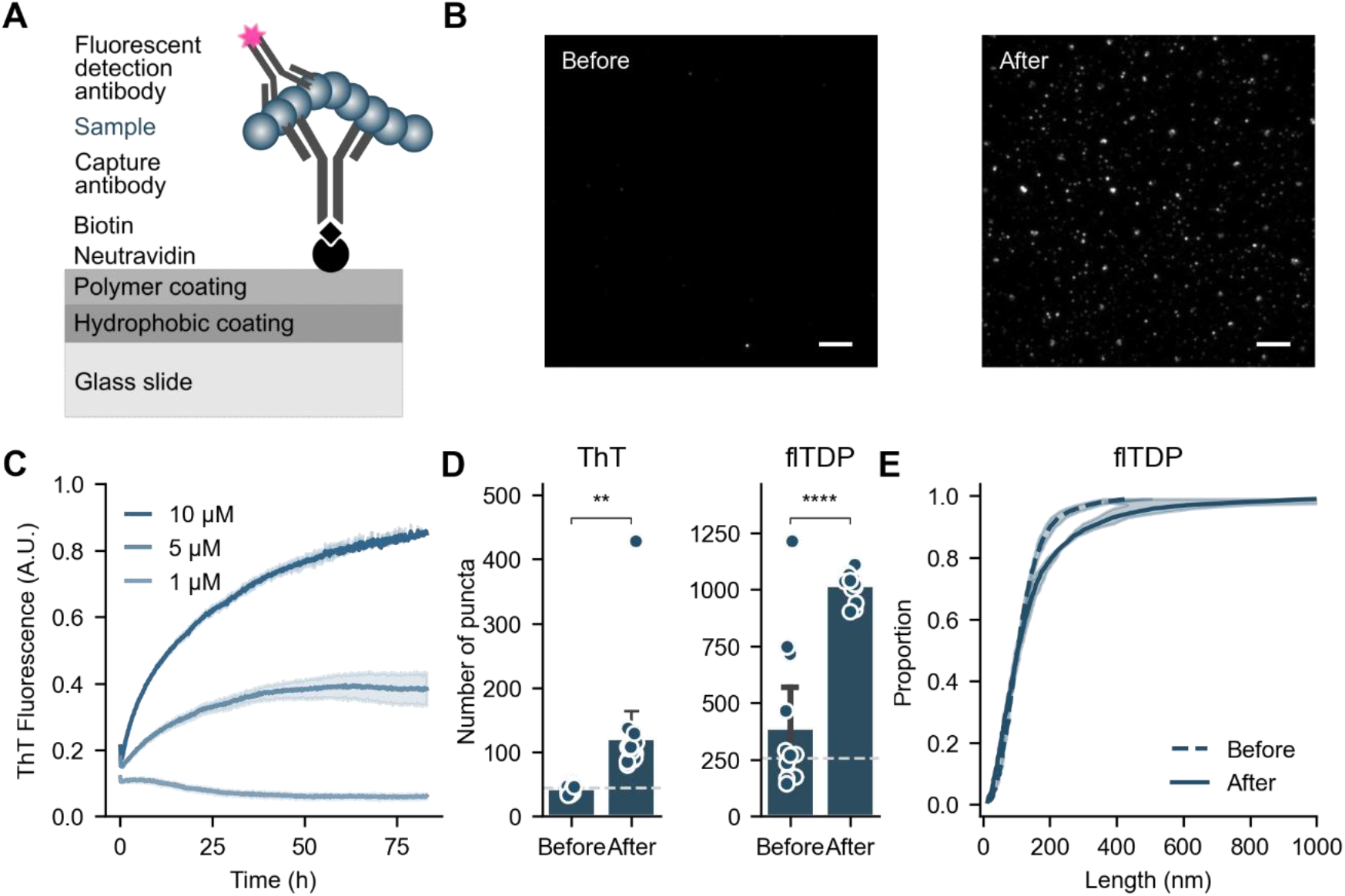
SiMPull provides a sensitive and specific method for detecting multimeric TDP-43 assemblies. **(A)** Schematic outlining the single-molecule pulldown (SiMPull) technology for detecting multimeric TDP-43 assemblies. Sequential assay steps are separated by washing, and the final fluorescent puncta quantified via TIRF microscopy. Numerous readouts are available from the assay, including diffraction-limited count and composition, and super-resolved size and shape. **(B)** Exemplar images of SiMPull performed before (0 h) and after (72 h) aggregation of the recombinant RRMs fragment. Scale bars indicate 5 µm. **(C)** Aggregation of RRMs ranging in concentration from 1 -10 µM was monitored via thioflavin-T (ThT) fluorescence at 490 nm over the course of 80 h. Shown are mean ± S.D. for three replicates. **(D)** SiMPull analysis of samples collected from 5 µM RRM solution before (0 h) and after (72 h) aggregation, as depicted in *B/C*. Diffraction-limited particle count is shown alongside a buffer-only control (dashed line), compared using a Welch’s t-test, ** *p* < 0.01, **** *p* < 0.0001. **(E)** Cumulative distribution of super-resolved particle length in samples collected before (0 h) and after (72 h) aggregation depicted in *B*. Shown in *D* are the mean ± S.D. of up to 16 fields-of-view, overlayed with the raw datapoints. Data are representative of three biological replicates.

We confirmed the selectivity of the TDP-43 SiMPull configuration for multimers over monomer (Figure 1B) using a recombinant aggregation-prone fragment of TDP-43 comprising the RRM1 and RRM2 domains^18^. As with other amyloid formation, concentration-dependent aggregation of this fragment can be monitored via thioflavin-T fluorescence^18,19^ (Figure 1C). Using aliquots of the protein solution collected immediately prior to (“Before”, 0 h) and following (“After”, 72 h) aggregation, we examined the samples via flTDP-SiMPull (Figure 1B-E). We found that, qualitatively, there was a large increase in the number of fluorescent puncta visible after aggregation (Figure 1B). Correspondingly, quantifying the number of puncta using either the thioflavin-T or flTDP antibody fluorescence revealed significantly more assemblies were present in the sample after aggregation (Figure 1D; *t* = −3.37 and −7.75, *p* = 0.0023 and *p* < 0.0001 respectively). Finally, we determined the size distribution of the captured assemblies before and after aggregation. Owing to the diffraction-limited nature of optical microscopy, we performed direct stochastic optical reconstruction microscopy (dSTORM), a super-resolution technique which facilitates accurate determination of aggregate length with 20 nm resolution^20^. We measured more than 3500 individual aggregates from samples taken before and after aggregation, finding that the samples after aggregation contained a higher proportion of longer aggregates (**Figure 1E;** *ks* = 0.110, *p* < 0.0001). Together, these data confirm the chosen SiMPull configuration provides a sensitive and specific method to capture and characterise multimeric TDP-43 assemblies.

### TDP-43 positive assemblies are not unique to disease

To assess the multimeric TDP-43 assemblies present in disease-derived brain tissues, we obtained a cohort of 10 individuals consisting of 5 MND donors with observed TDP-43 pathology (MND_*TDP*_) and 5 donors negative for TDP-43 pathology as a control cohort (Supplementary Figure 1). The control cohort included 3 neurologically normal donors and 2 disease-control donors with diagnosed familial SOD1-MND. Pathological aggregation of SOD1 and TDP-43 are generally considered to be mutually exclusive^1,21^. Intriguingly, however, our results suggested otherwise. Indeed, segregation of the SOD1-MND phenotype from the other control samples early in our analyses led us to separate the cohort into neurologically normal (CRL) and SOD1-MND (MND_*SOD*_) for all further analyses. For completeness, these results are presented here in the absence of any statistical comparisons owing to the small number of donor replicates. For each donor, we obtained matched frontal cortex (characterised by advanced TDP-43 pathology in MND) and cerebellar (which does not typically display TDP-43 pathology) tissue for analysis, together forming a set of 20 samples for characterisation. Each tissue sample was first soaked in an artificial CSF medium before the remaining tissue was homogenised (Supplementary Figure 1). This approach allowed us to characterise both diffusible protein assemblies, which may contribute to propagation to neighbouring cells or through the body, and embedded assemblies normally retained within the tissue, providing insight into the distribution and physicochemical properties of these subpopulations. The resultant extracts were characterised using SiMPull (Figure 2A; Supplementary Figure 2).

**Figure 2:**
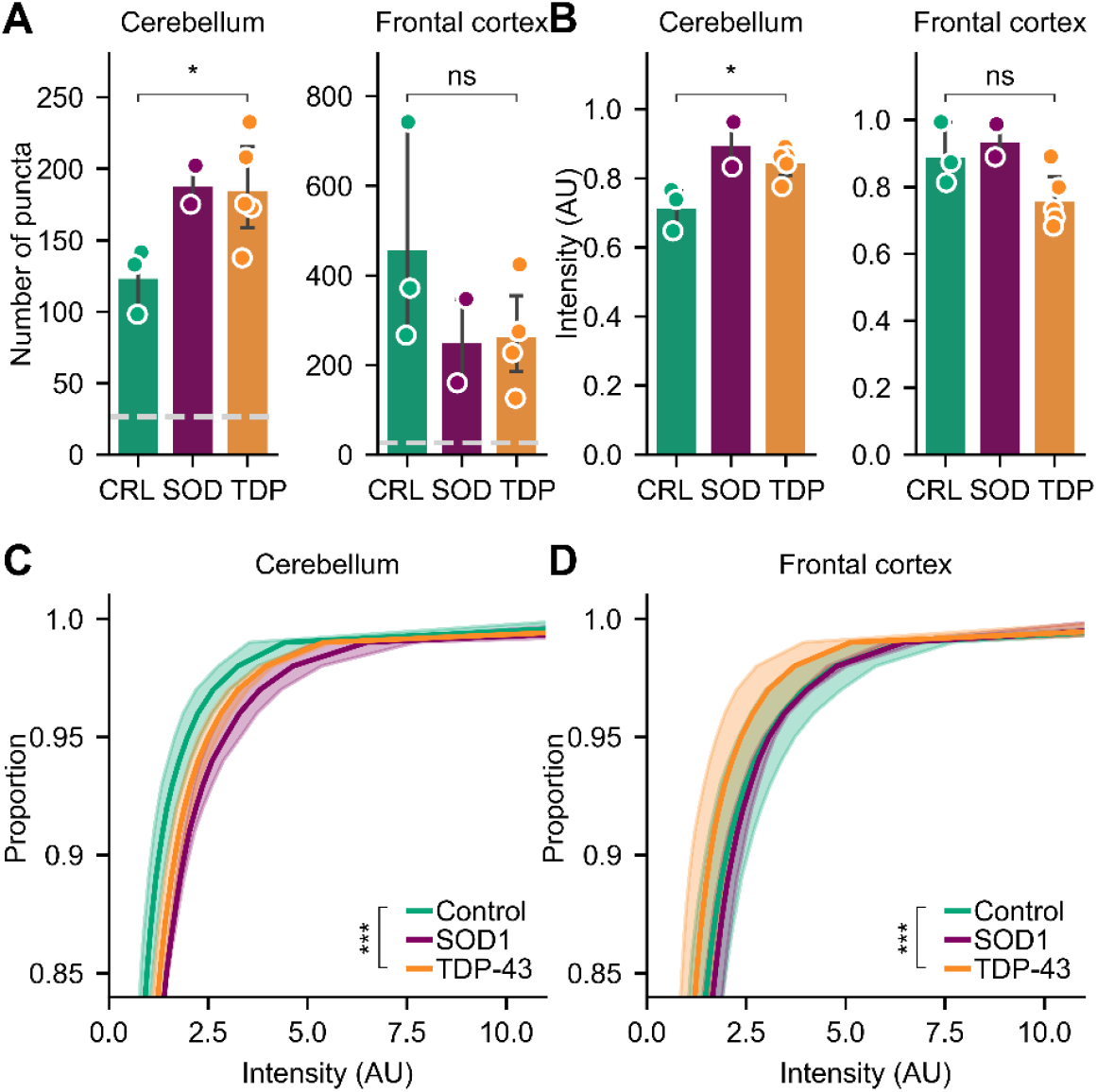
TDP-43 assemblies are found in both neurologically normal and disease-derived brain extracts. **(A)** Number and **(B-D)** brightness of diffusible TDP-43 positive assemblies captured with flTDP-SiMPull. Shown in *A, B* are the mean ± S.D. of donors in each cohort overlayed with the raw mean datapoints from three technical replicates for each donor, compared using Welch’s t-test, * *p* < 0.05, *ns p* > 0.05. Dashed line corresponds to an isotype control exposed to a reference sample of the same extract type. Shown in *C, D* are the mean interpolated cumulative distributions for each cohort ± S.D. of individual donors, compared via Kolmogorov–Smirnov test, *** *p* < 0.001.

Diffraction-limited SiMPull detected TDP-43 positive assemblies in all three of the cohorts tested, in both the diffusible and embedded extracts (Figure 2, Supplementary Figure 2). When samples were depleted of aggregates using an aggregate-specific chemical antibody (CAP1^22^), the number of TDP-43 positive assemblies detected via SiMPull was reduced to background, suggesting these assemblies are primarily aggregates in nature (Supplementary Figure 3). Intriguingly, significantly more TDP-43 positive assemblies were identified in the MND_*TDP*_ derived cerebellum extracts (Figure 2A; *t* = −2.57, *p* = 0.042). In contrast, MND_*TDP*_ derived frontal cortex contained fewer TDP-43 positive assemblies when compared to neurologically normal controls (Figure 2A; *t* = 1.58, *p* = 0.17). This phenomenon was mirrored when the flTDP capture antibody was replaced with CAP1 for SiMPull (Supplementary Figure 3), again supporting the assertion that the assemblies captured from tissue extracts are predominantly soluble nanoscopic aggregates. In addition to abundance, we analysed the intensity of individual puncta as a pseudo-measure of aggregate size (Figure 2C-D). While there was no significant difference in the mean intensity of assemblies between the control and MND_*TDP*_ cohorts in the frontal cortex extracts (Figure 2B; *t* = 2.07, *p* = 0.084), exploiting the single-molecule nature of SiMPull allowed us to detect significant differences in the distribution of intensities (Figure 2C-D; *ks* = 0.201 and 0.162 for cerebellum and frontal cortex respectively, *p* << 0.001 for both). This analysis revealed more, brighter TDP-43 assemblies in the cerebellum of MND_*TDP*_, while this trend was reversed in the frontal cortex. Together, these data support subtle but significant differences in disease-derived multimeric TDP-43 assemblies.

### Morphology of multimeric TDP-43 assemblies in disease

Super-resolution methodologies capable of resolving assemblies below the diffraction limit, such as dSTORM^20^, offer a way to determine particle morphology at the nanoscale. Using dSTORM, we conducted a detailed inventory of the morphology of individual TDP-43 positive assemblies captured using SiMPull. Exemplar images of individual assemblies with and without super-resolution reconstruction are shown (Figure 3A). These properties were then compared across each cohort.

**Figure 3:**
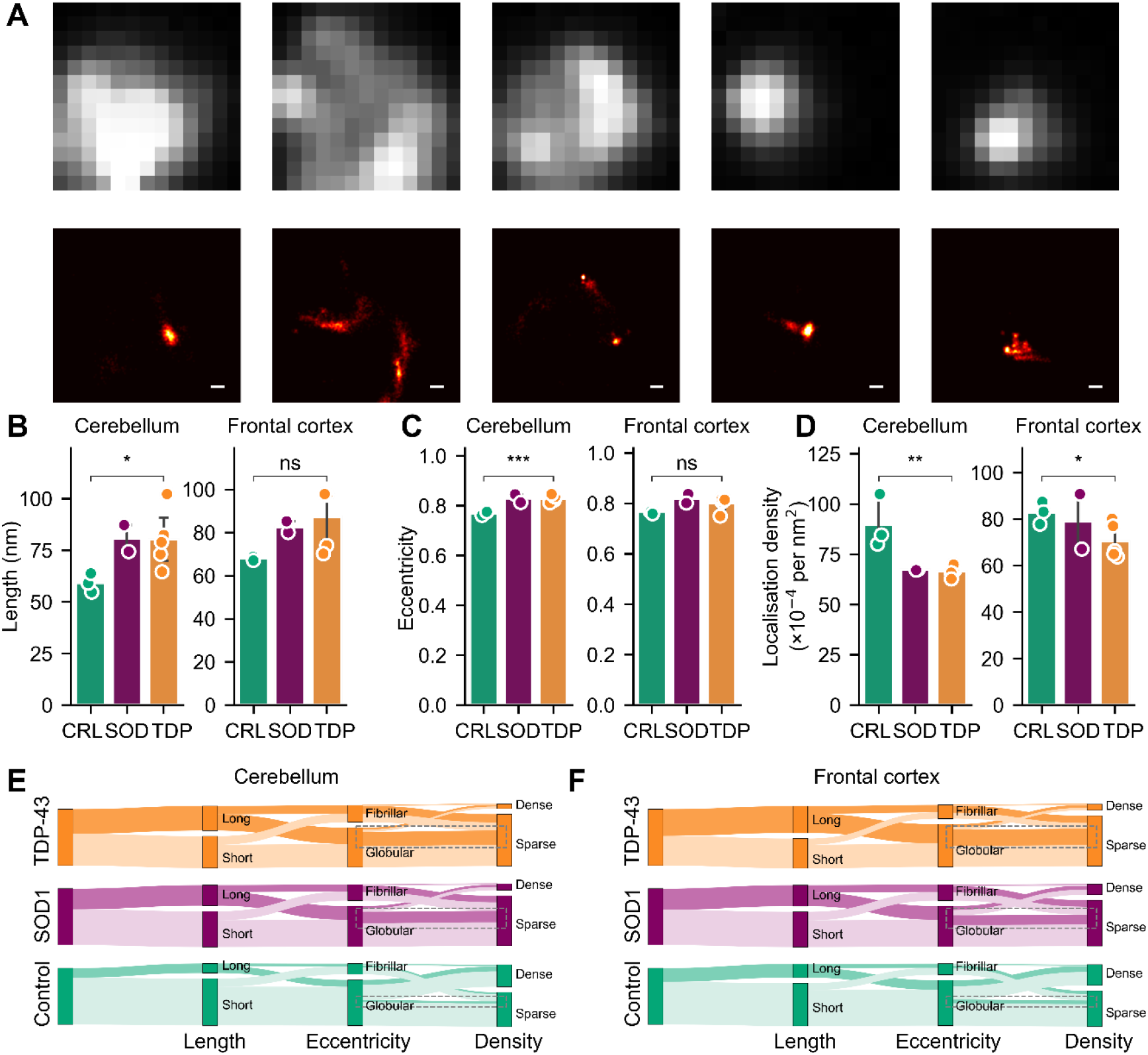
TDP-43-positive assemblies differ in size and shape between neurologically normal and disease-derived brain extracts. **(A)** Exemplar diffraction-limited (*upper*) and super-resolved (*lower*) regions of interest rendered from dSTORM of diffusible TDP-43-labelled assemblies. The same total area is shown in both upper and lower panels, and scale bars represent 100 nm. **(B-D)** Average **(B)** length, **(C)** eccentricity, and **(D)** localisation density of diffusible TDP-43 positive assemblies captured with CAP1-SiMPull. **(E-F)** Sankey visualisation of the proportion of assemblies that fall into each category as defined by cumulative thresholds for length (long > 100 nm), eccentricity (fibrillar > 0.9) and localisation density (dense > 0.01 localisations per nm^2^). Shown in *B*-*D* are the mean ± S.D. of donors in each category overlayed with the raw mean datapoints from three technical replicates of each donor, compared using Welch’s t-test, * *p* < 0.05, ** *p* < 0.01. Shown in *E, F* are the compiled assembly populations for each cohort. Dashed boxes indicate subpopulation of interest in all three cohorts.

We identified significant differences in the average of three morphological characteristics between cohorts; length, eccentricity and localisation density (Figure 3B-D; Supplementary Figure 4). Both length and eccentricity concern the overall shape of the assemblies, while localisation density is a measure of the distribution of accessible target epitopes within individual assemblies. MND_*TDP*_ derived TDP-43 positive assemblies tended to be longer (Figure 3B) and more elongated (Figure 3C), with these differences reaching statistical significance in the cerebellum-derived extracts (*t* = −2.46 and −6.77, *p* = 0.049 and 0.00051 respectively). In addition, the density of flTDP localisations per assembly was significantly reduced in disease-derived extracts from both the cerebellum and frontal cortex (Figure 3D; *t* = 4.03 and 2.50, *p* = 0.0069 and 0.047 for cerebellum and frontal cortex respectively). This difference supports the assertion that MND is associated with altered composition and/or structure of TDP-43 assemblies.

In addition to differences in the mean characteristics, distinct subpopulations of assemblies defined by their morphological signature were found to differ in prevalence in the MND_*TDP*_ derived extracts (Figure 3E-F). Exploiting the single-particle nature of SiMPull, it is possible to segregate individual assemblies according to their length (where ‘long’ ≥ 100 nm)^23^, eccentricity (where ‘fibrillar’ ≥ 0.9) and localisation density (where ‘dense’ ≥ 0.01 loc/*nm*^2^). A Sankey diagram then facilitates simultaneous visualisation of the proportion of all assemblies falling into each category and how distinct subpopulations characterised by membership of each category change between disease and neurologically normal brains (Figure 3E-F). We observed an increase in the proportion of long, fibrillar, sparsely labelled assemblies associated with disease in both the cerebellum and frontal cortex samples, mirroring the conclusions drawn from the mean comparison. In addition, this visualisation revealed a subpopulation of elongated, globular assemblies which were sparsely labelled that were increased in disease-derived extracts (Figure 3E-F; dotted outlines). These observations highlight the utility of single-particle measurements in detecting distinct subpopulations of TDP-43 containing assemblies that typify MND but are invisible to traditional methods or summary measures.

### MND-derived TDP-43 assemblies contain unique morphological signatures that are disease and tissue specific

The six parameters reported for TDP-43 positive assemblies so far support disease-associated differences (Figure 4A). Intriguingly, these differences were most pronounced and robust in the cerebellum samples where particle number, and the proportion of assemblies classed as long, fibrillar, and dense, were distinct between cohorts. However, owing to the single particle nature of SiMPull we can collect more than 15 parameters for every individual aggregate. Given the subtle differences observed in subpopulations between cohorts (Figure 3, Figure 4A), we reasoned a combination of all the parameters measured would most readily distinguish these cohorts. For this purpose, we chose a supervised dimensionality reduction tool, linear discriminant analysis (LDA), which ascertains the linear combination of parameters that best characterize or separate multiple categories.

**Figure 4:**
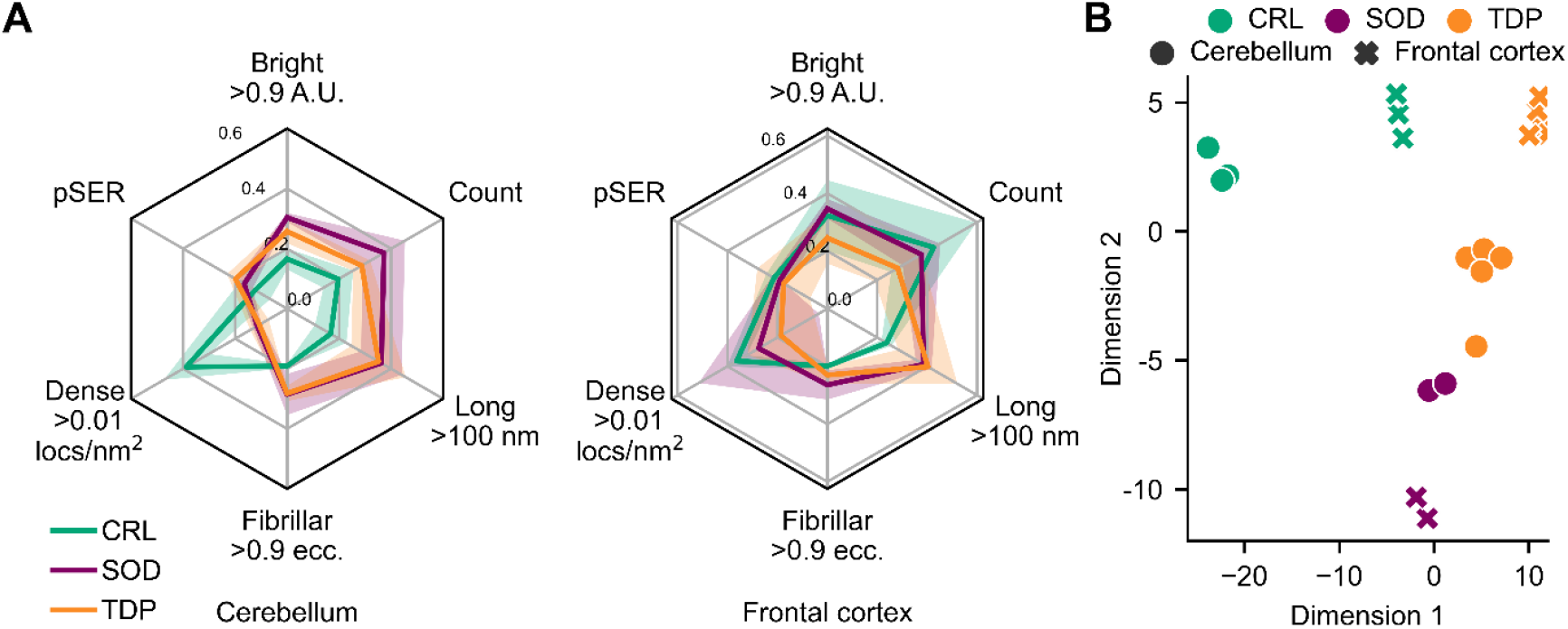
Signatures of MND found in nanoscopic TDP-43 assemblies. **(A)** Radar visualisation summarising the properties derived from diffusible samples characterised via SiMPull. Shown is mean ± S.D. for each cohort (CRL *n*=3, SOD1 *n*=2, TDP-43 *n*=5) across six measures; number of puncta as a proportion of the maximum count in any individual donor (count), proportion colabelled with pSER (pSER), proportion with length > 100 nm (long), proportion with eccentricity > 0.9 (fibrillar), proportion with localisation density > 0.01/nm^2^ (dense), and proportion whose intensity >0.9 A.U. (bright). **(B)** Two-dimensional projection of linear discriminant analysis performed on properties derived from diffusible samples characterised via SiMPull. For full list of included parameters, see Materials and Methods.

Linear discriminant analysis visibly separated each of the cohorts when projected onto a two-dimensional plane, as well as readily distinguishing extracts from different tissues (Figure 4B). Dimensions 1 and 2 encompassed 80.2% and 11.5% of the variance between categories respectively. Unsurprisingly, analysis of the resultant pseudo-eigenvectors ranks density, eccentricity, and length as the strongest contributors to this separation. This separation was equally evident when combining measurements from diffusible extracts with those from embedded extracts, and indeed more so when considering the embedded extract measurements alone for which the first two dimensions encompass 97.7% and 1.5% of the variance respectively (Supplementary Figure 5).

### The composition of nanoscopic TDP-43 assemblies is disease-specific

Pathological post-translational modifications of TDP-43 such as phospho-serine at position(s) 409/410 are considered a key identifier of pathological microscopic aggregates^24^; these modifications are also detectable in nanoscopic TDP-43 assemblies. Using a second fluorescently labelled detection antibody, SiMPull enables low-throughput compositional analysis whereby the proportion of target assemblies co-labelled for a second target of interest can be determined (Figure 5A). As anticipated, using an antibody against phospho-serine 409/410 (pSER) we observed a small but significant increase in the number of colabelled TDP-43 assemblies in MND_*TDP*_ derived frontal cortex embedded extracts (Figure 5B; *t* = −2.71, *p* = 0.035).

**Figure 5:**
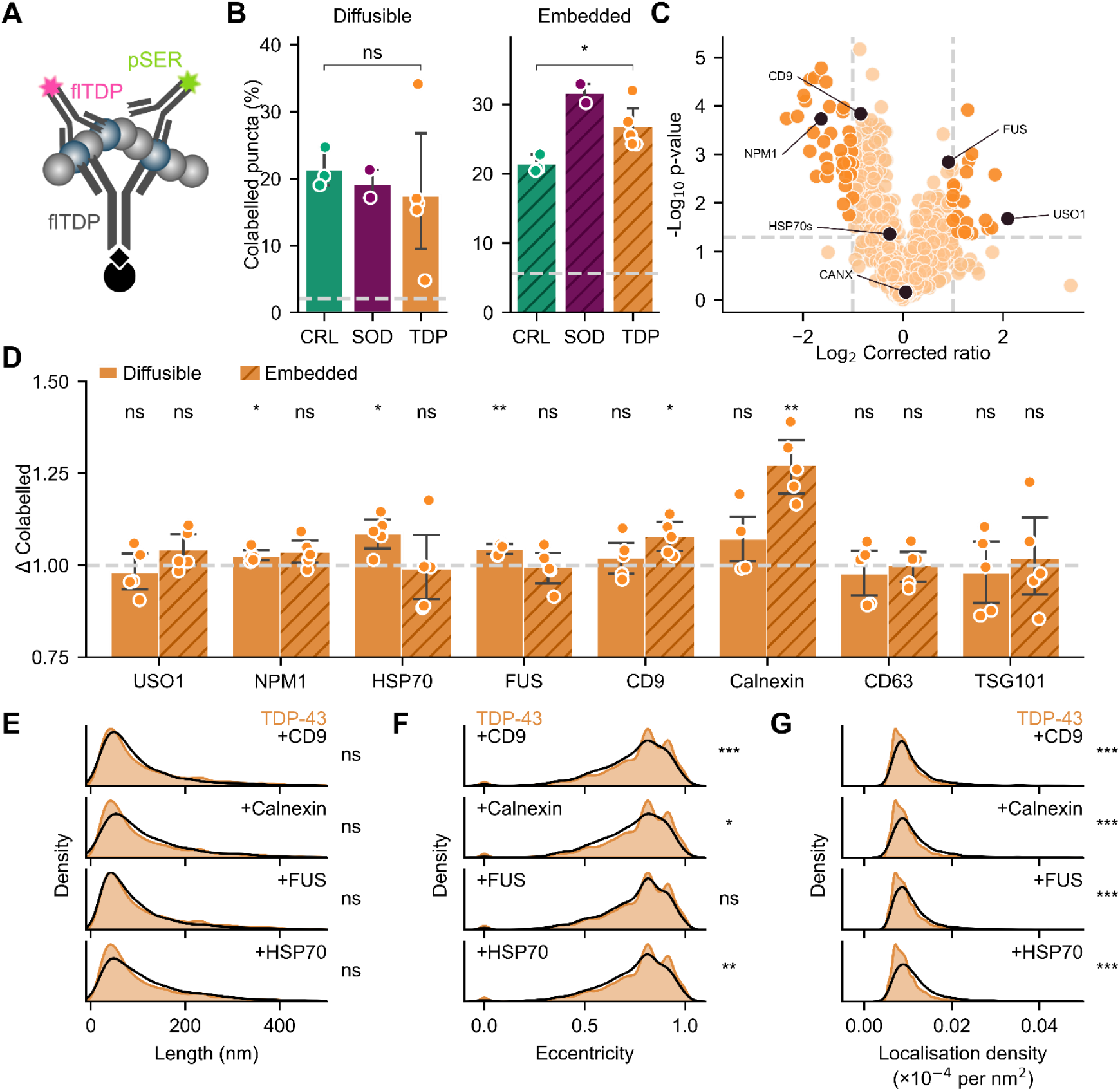
The composition of nanoscopic TDP-43 assemblies is altered in disease. **(A)** Schematic of the colabelling SiMPull configuration applied in *B*. **(B)** Proportion of TDP-43-positive assemblies taken from frontal cortex extracts also positive for pSER409/410. Shown are the mean ± S.D. of donors in each category overlayed with the raw mean datapoints from three technical replicates of each donor, compared using Welch’s t-test, * *p* < 0.05. Dashed line corresponds to an isotype control exposed to a reference sample of the same extract type. **(C)** Volcano plot depicting abundance ratio of proteins identified in the CAP1 aggregate fraction from frontal cortex embedded extracts of MND_*TDP*_ vs neurologically normal control cohorts. Dashed grey lines depict |*log*_2_*FC*| > 1 and −*log*_10_(*p*−*value*) > 1.3 thresholds outside which proteins are considered significantly different (dark orange). Proteins of interest (black) are highlighted. **(D)** Relative proportion of TDP-43 assemblies quantified via SiMPull positive for each protein of interest when compared to the control cohort (grey dashed line). Differences to control were assessed via one-sample t-test against the hypothetical mean of 1 with Bonferroni correction, * p < 0.05, ** p < 0.01. **(E-G)** Distribution of super-resolved **(E)** length, **(F)** eccentricity, and **(G)** localisation density for TDP-43 assemblies negative (orange) or positive (black) for various target proteins of interest. Distributions were compared via Kolmogorov–Smirnov test with Bonferroni correction. To avoid bias due to varying *n*, tests were bootstrapped using matched-*n* subsamples taken randomly from each population without replacement. * *p* < 0.05, ** *p* < 0.01, *** *p* < 0.001. Shown in *B, D* are the mean ± S.D. of donors in each category overlayed with the raw mean datapoints from three technical replicates of each donor.

To broaden our characterisation of TDP-43 assembly composition, we first identified proteins found in the aggregate fraction captured using the protein agnostic CAP1 for affinity purification and proteomics. Aggregates were isolated from each of the donor embedded extracts, subjected to isobaric labelling via tandem mass tags (TMT) and quantified alongside an equivalent total proteome sample via LC-MS/MS. To remove any bias resulting from differences in the total proteome composition, we first corrected the abundance ratio of proteins identified in the aggregate fraction according to the total proteome ratio of the same protein (Supplementary Figure 6A-B). The resultant corrected ratios were then partitioned using standard thresholds (|*log*_2_*FC*| > 1 and −*log*_10_(*p*−*value*) > 1.3) to yield the final list of proteins that were over-or under-represented in the aggregate fraction of MND_*TDP*_ derived extracts.

A total of 21 proteins were significantly enriched in the aggregate fraction, while 38 were significantly less abundant (Figure 5C) after correction for proteome-wide changes in abundance associated with disease. Gene ontology enrichment analysis using PantherGOslim identified no enriched ontology terms among these proteins, unlike the differences observed in the whole proteome (Supplementary Figure 6C). However, the collection of proteins over-represented in the aggregate fraction in MND_*TDP*_ derived extracts was significantly more connected by known protein-protein interactions than would be expected for similarly sized networks by chance (Supplementary Figure 6D, STRINGdb *p* = 0.0263). This is consistent with functionally linked protein networks being coprecipitated within aggregates^25^.

Target proteins were selected for evaluation via SiMPull, including NPM1 which was less abundant in the disease-derived aggregate fraction and USO1 which was more abundant in the disease-derived aggregate fraction (Figure 5C; black circles). In addition, we selected a panel of proteins commonly associated with exosomes; CD9, CANX, and HSP70 for which disease-associated differences were considered not significant (Figure 5C; black circles), and TSG101 and CD63 which were not identified in our proteomic analyses. Secretion in extracellular vesicles has been identified as a key pathway for clearance of TDP-43 from the cytoplasm^26^ which, when inhibited, results in the accumulation of cytoplasmic TDP-43 foci^27^. HSP70 shows robust recruitment to such cytoplasmic TDP-43 foci in cellular models^28^. Finally, we selected FUS, a protein similarly associated with MND^29^ and which has been found to form insoluble aggregates in cell models overexpressing TDP-43^30^. Of these, TDP-43 shares known physiological interactions with FUS (high confidence; score = 0.985), NPM1 (medium confidence; score = 0.376) and CANX (low confidence; score = 0.285) (STRINGdb). We assessed the association of each of these proteins with aggregates containing TDP-43 using the CAP1-SiMPull configuration.

Significantly more TDP-43 assemblies contained NPM1, HSP70, FUS, CD9 and CANX in the MND_*TDP*_ cohort when compared to neurologically normal controls (Figure 5D). This is despite no difference in the total number of target-positive puncta (Supplementary Figure 7), and a corresponding lack of significant difference in the total abundance of HSP70, FUS, CD9 and CANX in the aggregate fraction as measured via proteomics. Leveraging super-resolved morphological measurements of individual particles allowed us to separate the characteristics of those TDP-43 aggregates decorated with these target proteins from those without (Figure 5E-G). There was no bias in colabelling according to the length of TDP-43 assemblies (Figure 5E). However, significant differences in the distribution of eccentricity and localisation density associated with TDP-43 assemblies containing HSP70, FUS, CD9 and CANX were observed when compared to assemblies negative for these proteins. Colabelled assemblies lacked a more eccentric (Figure 5F), sparsely labelled (Figure 5G) subpopulation present in the singly labelled assemblies. This missing population is reminiscent of the features found to be associated with TDP-43 assemblies enriched in the MND_*TDP*_ cohort (Figure 3C-D). Together this suggests complex modulation of TDP-43 assemblies in disease; sparsely labelled fibrillar TDP-43 assemblies accumulate, however a population of globular, densely labelled TDP-43 assemblies is more frequently engaging cellular export machinery. These observations underscore the utility of single-molecule methods in characterising and untangling intricate signatures of MND.

## DISCUSSION

Single-molecule microscopy provides unparalleled resolution and sensitivity for the investigation of disease-associated protein aggregates^12,13^. Here, we applied two novel single-molecule pulldown configurations to quantify and characterise protein assemblies containing TDP-43. We observed less TDP-43-positive assemblies in MND_*TDP*_ frontal cortex extracts compared to neurologically normal controls, however those assemblies tended to be longer, more fibrillar, and sparsely decorated by antibodies targeting the RRMs motif. This is entirely consistent with unmasking of mature TDP-43 aggregates, where the C-terminal amyloid core of fibrillar aggregates is exposed by cleavage of the RRM domains^31^. Importantly, we show these characteristics are sufficient to distinguish extracts from disease and control cohorts. Given these differences are observed in the diffusible as well as embedded extracts, they are highly likely to be evident in more accessible biofluids^32,33^ positioning this technology as a potent potential diagnostic and disease staging tool.

That the relative abundance of MND_*TDP*_ -derived TDP-43 positive assemblies was lower in the frontal cortex but higher in the cerebellum is counterintuitive. However, unmasking of end-stage aggregates is likely to be most prominent in the frontal cortex as this region is affected early in disease^34^, allowing aggregates to mature. Combined with the higher relative abundance of pSER-positive assemblies in the frontal cortex, these results are consistent with the frontal cortex exhibiting advanced TDP-43 aggregate pathology that has proceeded over the entire disease course. In contrast, spreading of TDP-43 pathology through the brain^34,35^ reaches the cerebellum late in the disease course meaning we would anticipate immature TDP-43 aggregates to predominate at the end stage of disease. Indeed, cerebellar TDP-43 pathology was variably detected using classical immunohistochemistry approaches^36^, and has only recently been reliably detected through the use of sensitive molecular technologies^36,37^. Consistent with this, we found more cerebellum-derived TDP-43 positive assemblies in MND_*TDP*_ and no significant difference in pSER positivity. Thus, these assemblies likely reflect early-stage aggregates that have not yet been unmasked.

Our exploration of aggregate composition revealed additional subtle differences between disease and control extracts. The composition of disease-derived aggregates offers insight into disease mechanisms by identifying potentially toxic loss-or gain-of-function among proteins found in aggregates. Recently, proteomic quantification of insoluble aggregate fractions from various disease models and tissues has delivered several such datasets^38^. However, our unique combination of proteomic and microscopic methods highlights the necessity of considering heterogeneity in aggregate composition, a property which is inaccessible to bulk proteomic methods. Of the examples we examined, which included proteins with a range of relative enrichment in the aggregate fraction, we found little correlation between the relative abundance of the protein in the aggregate fraction and the likelihood of these proteins being associated with individual TDP-43 positive aggregates. Instead, we observed subpopulations of aggregates whose morphology differed according to whether these proteins were present in the aggregate. Importantly, the subpopulation of TDP-43 positive aggregates negative for additional proteins matched the morphology of the dominant subpopulation associated with disease, i.e. they tended to be more fibrillar and sparsely decorated by the RRMs motif. These results provide important context for interpreting proteomic studies and highlight the complexity of targeting coaggregating proteins as an approach for therapeutic intervention.

Differences in the MND_*SOD*_ and MND_*TDP*_ cohorts, while not a focus of this study, were unexpected. Traditionally, MND_*TDP*_ and MND_*SOD*_ have been considered mutually exclusive when it comes to the presence of TDP-43 aggregates^1,21^. However, increasingly sensitive methods now allow us to quantify the early stages of TDP-43 dysfunction and aggregation. Indeed, two additional orthogonal methods (Supplementary Figure 8; Supplementary Figure 9; Supplementary Table 1) showed similar evidence of TDP-43 dysfunction in both the MND_*SOD*_ and MND_*TDP*_ cohorts when compared to neurologically normal controls. This was not generic; the trend of similarity between MND_*SOD*_ and MND_*TDP*_ was not observed in the abundance of Apoptosis-associated Speck-like protein containing a CARD (ASC) specks^39^, an indicator of inflammasome activation (Supplementary Figure 10). While the small sample size prevents us from making statistically driven conclusions, this will be a priority for future studies exploring the presence and role of TDP-43 dysfunction in MND_*SOD*_ with this new generation of sensitive molecular technologies.

Single molecule methods provide unprecedented detail in our effort to understand TDP-43 aggregation in the context of neurodegenerative disease, including MND. Understanding the complex interplay between genetic and environmental factors in the formation and propagation of physiological and pathological TDP-43 assemblies is vital for therapeutic strategies aimed at halting or mitigating the progression of disease-associated TDP-43 aggregation. Here we demonstrate that quantifying aggregate composition and morphology via SiMPull yields aggregate characteristics with sufficient detail to resolve disease cohorts and tissues from their neurologically normal counterparts. We believe such technologies are the first step toward next-generation diagnostic approaches, requiring small sample volumes and minimally invasive sample collection methodologies.

## MATERIALS AND METHODS

### Materials

Purified recombinant TDP-43 RRM constructs as described by Zacco and colleagues^18^ were kind gifts of Dr Alexandra Louka and Dr Annalisa Pastore. Antibodies for TDP-43 (ProteinTech #60019-2-Ig), pSER409/410 (ProteinTech #66318-1-Ig), calnexin (AbCam #ab133615), CD9 (AbCam #ab263019), CD63 (AbCam #ab134045), Hsp70 (AbCam #ab181606), TSG101(AbCam #ab125011), and nucleophosmin (ProteinTech #60096-1-Ig) were labelled in house using the biotinylation (AbCam #ab201796) or AlexaFluor (AbCam #ab269823) conjugation kits provided by AbCam according to the manufacturer’s instructions. The resultant conjugates were then purified (AbCam #ab102778) before being quantified and stored at 4°C until use. The CAP1 reagent was a generous gift of Dr Margarida Rodrigues (University of Cambridge), provided as 2 mM stocks in DMSO stored at −20 °C until use.

### Tissue details

Freshly frozen post-mortem brain samples were provided by the London Neurodegenerative Diseases Brain Bank (KCL), part of the UK Brain Banks Network. The project was carried out under the ethical approval of the tissue bank (18/WA/0206 REC for Wales), who were responsible for obtaining written informed consent from the donors and/or their relatives as appropriate. Samples included matched frontal cortex (BA) and cerebellum samples taken from 5 donors with TDP-43 pathology. These donors were compared to an age-matched control cohort lacking TDP-43 pathology, consisting of 2 donors diagnosed with SOD1-MND and 3 neurologically normal donors. Donor characteristics are summarised in Table 1, for which no significant differences were identified between each cohort.

**Table 1:** Tissue donor characteristics.

| **MRC BNN-ID** | **Category** | **Sex** | **Age** | **Post-mortem delay (min)** |
| --- | --- | --- | --- | --- |
| **BBN_6209** | SOD1-MND | F | 70 | 56 |
| **BBN_16553** | SOD1-MND | F | 46 | 5 |
| **BBN002.28672** | TDP43-MND | F | 54 | 67.5 |
| **BBN_24296** | TDP43-MND | M | 68 | 73 |
| **BBN_11078** | TDP43-MND | M | 61 | 20.5 |
| **BBN_6280** | TDP43-MND | M | 75 | 38 |
| **BBN002.28946** | TDP43-MND | F | 49 | 64 |
| **BBN002.33764** | Control | M | 73 | 49 |
| **BBN002.30139** | Control | F | 47 | 27 |
| **BBN_21801** | Control | M | 68 | 60 |

### Preparation of human brain extracts

Brains were flash-frozen and stored at −80°C at the London Neurodegenerative Disease Brain Bank. Extracts from each donor were prepared as described previously^40^. Briefly, approximately 120 mg samples were dissected and transferred to 5 volumes of ice-cold buffer mimicking cerebrospinal fluid (artificial CSF; 124 mM NaCl, 2.8 mM KCl, 1.25 mM NaH2PO4, 26 mM NaHCO3, pH 7.3 containing protease inhibitors 5 mM ethylenediaminetetraacetic acid (EDTA), 1 mM ethyleneglycoltetraacetic acid, 5 μg/ml leupeptin, 5 μg/ml aprotinin, 2 μg/ml pepstatin, 120 μg/ml Pefabloc and 5 mM NaF). Tissue was incubated for 2 h with gentle side-to-side mixing before being centrifuged at 2000×*g* for 10 min at 4°C. The upper 90% of the supernatant was removed and centrifuged at 21000×*g* at 4°C for 60 min. The resulting supernatant was collected and dialysed against a 100-fold excess of fresh artificial CSF lacking protease inhibitors using Slide-A-Lyzer™ G2 Dialysis Cassettes (2K MWCO, Fisher Scientific) at 4 °C, with buffer changed three times over a 72 h period. Finally, the dialysed material was collected, designated as the diffusible extract, and aliquoted before being immediately snap frozen and stored at −80 °C before use.

Embedded extracts were prepared via bead-based homogenisation of the pellets generated using a method adapted from Goedert and colleagues^41^. Briefly, the 2000 ×*g* pellet was resuspended in 10 volumes of ice-cold homogenisation buffer (10 mM Tris-HCl, 0.8 M NaCl, 1 mM EGTA, 0.1% Sarkosyl, 10% sucrose; pH 7.32 containing cOmplete Ultra Protease Inhibitor and PhosStopTM Phosphatase Inhibitor) and transferred to tubes containing 1.5 mm triple-pure high impact zirconium beads (Benchmark Scientific). Samples were homogenized at 4°C in a VelociRuptor V2 Microtube Homogenizer (Scientific Laboratory Supplies) using two cycles of 20 sec shaking at 5 meters/second, with a 10-second rest between cycles. The resulting homogenate was centrifuged at 21000×*g* for 20 min at 4 °C, and the upper 90% of the supernatant was retained. This process was repeated with the resulting pellet using 5 volumes of homogenisation buffer, then the upper 90% of the supernatant was again removed. Both supernatant aliquots were combined, and this mixture was designated the embedded extract. Each sample was aliquoted before being immediately snap frozen and stored at −80 °C before use.

The total protein concentration for both diffusible and embedded extracts was determined using a bicinchoninic assay (Thermo Fisher) as per the manufacturer’s instructions. Optionally, the TDP-43 concentration was determined using a human TDP-43 ELISA kit (ProteinTech) according to the manufacturer’s instructions.

### Aggregation of RRM domains

The aggregation of recombinant RRM proteins was monitored using thioflavin-T (ThT) fluorescence as described previously^40^. Briefly, protein aliquots were rapidly thawed and centrifuged at 4 °C for 60 min at 100 000 ×*g* to remove any pre-formed oligomers. The concentration was then determined via spectroscopy to ensure equivalent starting concentrations across all assays. Protein aliquots were then diluted to a 2× stock in PBS before being combined 1:1 with high-salt phosphate buffer (20 mM potassium phosphate buffer pH 7.2, 300 mM KCl) containing thioflavin-T (20 µM) to arrive at the final assay concentration. Aliquots were transferred to a microplate in triplicate alongside buffer-only controls. The thioflavin-T fluorescence was then monitored at 440 nm / 490 nm excitation and emission respectively using a CLARIOstar plate reader (BMG Labtech) every 10 minutes for up to 80 hours. Following aggregation, aliquots to be used as positive controls in subsequent assays were collected and snap frozen before being stored at −80 °C until use.

### Single-molecule pulldown (SiMPull)

Single-molecule pulldowns specific for TDP-43 aggregates were adapted from previous versions targeting other proteins^12,15^ with the following modifications. All steps were completed at room temperature and all buffers filtered at 0.02 µm unless otherwise stated. Borosilicate glass coverslips (26 mm × 76 mm #1.5; VWR) were first cleaned using an argon plasma cleaner (PDC-002, Harrick Plasma) for 15 min to remove any fluorescent residues. One 50-well silicone gasket (Grace Bio-Labs, SKU 103250) was then attached to the surface and each well was coated with 6.5 μl of Rain-X Repellant for 2 min. Wells were washed three times with 10 µl ultrapure water then left to dry completely for up to 15 min.

Wells were first incubated with neutrAvidin at 0.2 mg/ml in PBS containing 0.05 % (v/v) Tween-20 (PBST) for 10 min before being washed three times with PBS. Wells were coated with 1 % (v/v) pluronic F-127 for 30 min followed by a further three washes with PBS, then incubated for 5 min with PBS containing 1 % (v/v) Tween-20. Biotinylated capture antibody or capture molecule for amyloid precipitation (CAP1) were prepared at 2 nM or 5 nM in PBST. Wells were incubated with capture solution for 30 min before being washed three times with PBST, with the final wash solution left to incubate for 10 min before continuing. Samples were then added to individual wells and incubated for 1 h in a humid chamber. Wells were again washed three times with PBST, then incubated with fluorescent detection reagents prepared at a final concentration of 0.5 −2 nM in PBST for 30 min protected from light. Wells were finally washed three times in PBST. For diffraction-limited imaging, wells were filled with fresh PBS before the gasket was sealed with another clean coverslip. For imaging to be super-resolved, wells were filled with freshly prepared dSTORM buffer (1 mg/ml glucose oxidase, 52 µg/ml catalase and 50 mM MEA (cysteamine) in 50 mM Tris in PBS + 10% (v/v) glucose, pH 8).

### Total Internal reflection Fluorescence (TIRF) microscopy

Fluorescent images were collected using a home-built total internal reflection fluorescence (TIRF) microscope as described previously^12^, consisting of an inverted Ti-2 Eclipse microscope body (Nikon) fitted with a 1.49 N.A., 60x TIRF objective (Apo TIRF, Nikon) and a perfect focus system. Images were acquired using a 638 nm laser (Cobolt 06-MLD-638, HÜBNER), 561 nm laser (Cobolt 06-DPL-561, HÜBNER), 488 nm laser (Cobolt 06-MLD-488, HÜBNER) or 405 nm laser (Cobolt 06-MLD-405, HÜBNER). Detection antibodies labelled with AF647, AF594 or AF488 were excited at 638 nm, 561 nm and 488 nm respectively, while CAP1 was excited at 405 nm. The resultant fluorescence was collected by the objective, passed through a quad-band dichroic beam splitter and cleaned up using the appropriate emission filter (for 488-nm-induced fluorescence: BLP01-488R-25×36, Semrock and FF01-520/44-25×36, Semrock; for 638-nm-induced fluorescence BLP01-635R-25×36, Semrock). Images were collected on an EMCCD camera (Evolve 512, Photometrics) operating in frame-transfer mode (electron-multiplying gain of 6.3 electrons/ADU and 250 ADU/photon) where each pixel corresponds to a length of 107 nm on the recorded image, resulting in individual fields of view of 54.784 µm^2^. Images were collected in a grid using an automated script (Micro-Manager) to avoid any bias in the selection of fields of view.

For diffraction-limited imaging, lasers were typically set at 1 - 2 % of the maximum power. Up to 16 fields of view were imaged per well each for 50 frames of 50 ms exposure. For co-localization experiments images were acquired for each field of view sequentially in each excitation channel. For super-resolved imaging, lasers were set to maximum power (190 mW and 120 mW for the 638 and 561 nm respectively), with 4 fields of view typically imaged per well each for 5000 frames of 33 ms exposure.

### Diffraction-limited image analysis

Fluorescent puncta were automatically detected in diffraction-limited TIRF images using a python-based adaptation of ComDet, as described previously^12^. Briefly, a mean intensity projection was prepared from the last 40 frames of each field of view to effectively minimise background noise, then thresholds optimised using positive (RRMs) and negative (isotype or monomeric controls) were applied via ComDet to isolate individual fluorescent spots. Optionally, to facilitate comparisons across technical replicates, and between samples the number of fluorescent spots detected in each field of view was corrected by a scaling factor combining the original sample weight before processing and the total protein concentration of the sample being assayed. Finally, characteristics for individual spots such as the mean intensity were measured using scikit-image^42^.

For colabelling experiments, ComDet was applied independently to both detection channels and the Euclidean distance between pairs of coordinates in the resultant list of centroids was calculated using SciPy^43^. Spots co-labelled in the opposing channel were assigned as those within a 4-pixel radius according to the Euclidean distance. In the case where more than one spot met this criterion, the closest spot was selected. Finally, the number of spots co-labelled by chance was estimated by transposing the centroid coordinates for the second channel in the x dimension and repeating the threshold distance analysis. The proportion of co-labelled spots was calculated as the number of co-labelled spots divided by the total number of spots detected in that channel.

### Super-resolved image analysis

Super-resolution images were reconstructed using the Picasso^44^ package as described previously^12^. Briefly, after discarding up to the first 300 frames, localizations were identified and fit then corrected for microscope drift using the inbuilt implementation of redundant cross-correlation. Localizations were then filtered for precision <30 nm, with a final mean precision of » 12 nm. Random localizations were removed using DBSCAN as provided by the scikit-learn package^45^ with parameters a maximum radius of 0.5 pixels and a minimum density of 5 localisations. The resultant clustered array of localizations was then subjected to a series of morphological dilation, closing and erosion operations as provided by the scikit-image package^42^ to yield single connected regions of interest corresponding to individual assemblies. Aggregates were then measured using a combination of scikit-image for basic region properties such as perimeter, area, and eccentricity, and SKAN^46^ for skeletonized length whereby the length of each aggregate is reported as the summed branch distance. Finally, super-resolved images were rendered using the inbuilt Picasso functionality.

### Cumulative distribution interpolation

To combine cumulative distributions for individual samples, it is necessary to interpolate at discrete intervals along the proportional axis. This was achieved by first calculating the cumulative distribution for individual samples, then the value at a series of 100 equidistant increments from 0 to 1 (0, 0.01, 0.02, 0.03 … 0.99, 1) was then interpolated using the ‘nearest’ method provided by pandas^47^. For statistical comparison of the distributions, to avoid the confounding factor of population size, a subset of 1000 assemblies was sampled at random from each cohort and compared using a Kolmogorov–Smirnov test. In the event multiple comparisons were performed, the resultant p-values were adjusted via either Holm-Bonferroni or Bonferroni correction for per-cohort or per-population comparisons respectively.

### Disease signature analyses

Linear discriminant analysis (LDA) was performed using Python’s scikit-learn package^45^ with combined region-cohort classes (cerebellum-control, cerebellum-SOD-MND, cerebellum-TDP-43-MND, frontal cortex-control, frontal cortex-SOD-MND, and frontal cortex-TDP-43-MND). The input data was derived from the following collection of SiMPull parameters: mean assembly count, area, eccentricity, perimeter, minor axis length, major axis length, number of fluorescent localisations, smoothed length, localisation density, scaled fluorescence intensity, and the proportion of assemblies characterised as bright, colocalised with phospho-serine, fibrillar, long and sparse. The resulting LDA model was used to visualize separation of the categories in two dimensions. Pseudo-Eigen’s were also calculated as a measure of the relative contribution of parameter to the final class separation.

### Affinity precipitation and peptide preparation

Affinity precipitation of aggregates from embedded samples was completed as described previously^22^. Briefly, streptavidin Dynabeads (MyOne Streptavidin C1, Invitrogen) were conjugated with CAP1 by first removing 10 μl of beads/sample from the vial after brief mixing. The beads were diluted 1:100 with PBS and placed on a magnet for 2– 3 min before discarding the supernatant, and this wash process was repeated three times. To load beads with CAP1, washed beads were resuspended in 2 ml of CAP1 (30 μM) and placed in a revolving mixer to incubate at room temperature for 1 h. Following this, beads were separated from the supernatant by magnet for 2–3 min and the supernatant discarded. Finally, the beads were washed three times with PBS and were then ready for use.

To isolate aggregates, 150 μl aliquots of embedded extract was combined with 10 μl of prepared beads and the resulting suspension was placed in a revolving mixer to incubate at room temperature for 2 h. After incubation, beads were separated from the depleted supernatant by magnet for 2–3 min and the supernatant discarded. Beads were then washed twice with PBS to minimise non-specific background before proceeding to proteomic analysis.

### Proteomic analysis

Total protein was precipitated from embedded extracts by combining with 5× ice-cold acetone and incubating overnight at −20 °C. Precipitated proteins were then isolated via centrifugation at 20000×g for 30 min at 4 °C. Total protein and affinity-captured aggregates were then prepared alongside pooled reference samples for mass spectrometry using iST-NHS (Preomics) and tandem mass tag 11-plex (ThermoFisher Scientific) kits according to the manufacturer’s instructions. The resulting lyophilized peptides were quantified via bicinchoninic acid (BCA) assay (ThermoFisher Scientific) and the final concentration of peptides adjusted to 0.1 µg/µl in LC-LOAD buffer (Preomics).

Pooled fractions underwent LC–MS/MS analysis (Orbitrap Lumos Fusion Tribrid Mass spectrometer). Peptide samples were analysed on a Orbitrap Fusion Lumos Tribrid mass spectrometer interfaced with a 3000 RSLC nano liquid chromatography system. 1ug of each sample were loaded on to a 2 cm trapping column (PepMap C18 100A – 300 mm. Part number: 160454. Thermo Fisher Scientific) at 5 ml/min flow rate using a loading pump and analyzed on a 50 cm analytical column (EASY-Spray column, 50 cm 75 mm ID, Part number: ES803) at 300 nl/min flow rate that is interfaced to the mass spectrometer using Easy nLC source and electro sprayed directly into the mass spectrometer. A linear gradient between solvent A comprising of 0.1% Formic acid (Fisher) in water (Fischer) and solvent B comprising of 0.1% Formic Acid in 99.9% Acetonitrile (Fisher). Firstly a gradient of 3% to 30% of solvent-B was used at a 300 nl/min flow rate for 105 min and 30% to 40% solvent-B for 15 min and 35% to 99% solvent-B for 5 min which was maintained at 90% B for 10 min and washed the column at 3% solvent-B for another 10 min comprising a total of 140 min run with a 120 min gradient in a data dependent acquisition (DDA) mode. TMT tags were quantified using the SPS mode at MS3 level.

Initial peptide identification and quantitation was carried out using MaxQuant (version 2.6.2.0) against the Swissprot Homo Sapiens database (downloaded 2023.06.05; 20423 entries). The search was conducted with 20 ppm MS tolerance, 0.6 Da MS/MS tolerance, with one missed cleavage allowed and match between runs enabled. Variable modifications included methionine oxidation and N-terminal protein acetylation, while the PreOmics cysteine alkylation (+113.084Da) was set as a fixed modification. The false discovery rate maximum was set to 0.005% at the peptide identification level (the actual was 0.005 for each replicate) and 1% at the protein identification level. All other parameters were left as default.

Further analysis was performed with custom Python scripts^25,48^. First, the common contaminants including keratin were removed. Quantified proteins were considered as those identified by at least two peptides, one of which was unique. Protein abundances were corrected per-run (total) or per-channel (affinity-captured aggregates) based on the summed abundance of a reference sample. Abundance ratios were then calculated for each TDP-43-MND sample using the corresponding mean control abundance. Affinity-captured ratios were further normalised according to the total protein ratio. The resulting ratios were then subjected to a one-sample t-test against the hypothetical mean of 1, corresponding to no difference between disease and control. Finally, thresholds on the resultant mean abundance (|log_2_(abundance)| > 1) and p-value (-log_10_(*p*-value) > 1.3) were applied to determine proteins differentially abundant in TDP-43-MND.

### TDP-43 loss-of-function PCR splicing assay

For RNA extraction, brain tissue was thawed directly in TRIsure reagent (Bioline) and RNA isolated following manufacturer’s instructions. RNA was purified (RNeasy Plus Kit, Qiagen) and quantified using a Nanodrop Spectrophotometer (Thermo Fisher Scientific).

RNA (500 ng) was reverse transcribed (QuantiTect Reverse Transcription Kit, Qiagen) and the output volume of 20 μl diluted tenfold in nuclease free water (Promega). Real-time PCR was performed using specific primers (Merck; Table 2) and PowerUp SYBR Master Mix (Thermo Fisher Scientific) on a QuantStudio 6 Flex instrument using fast cycling mode based on the user guide (publication number MAN0013511 Rev. F.0, applied biosystems). Reaction specificity was confirmed by melt curve analysis and normalised expression (ΔΔCq) calculated manually using three reference genes *GAPDH, ACTB* and *YWHAZ*.

**Table 2:**
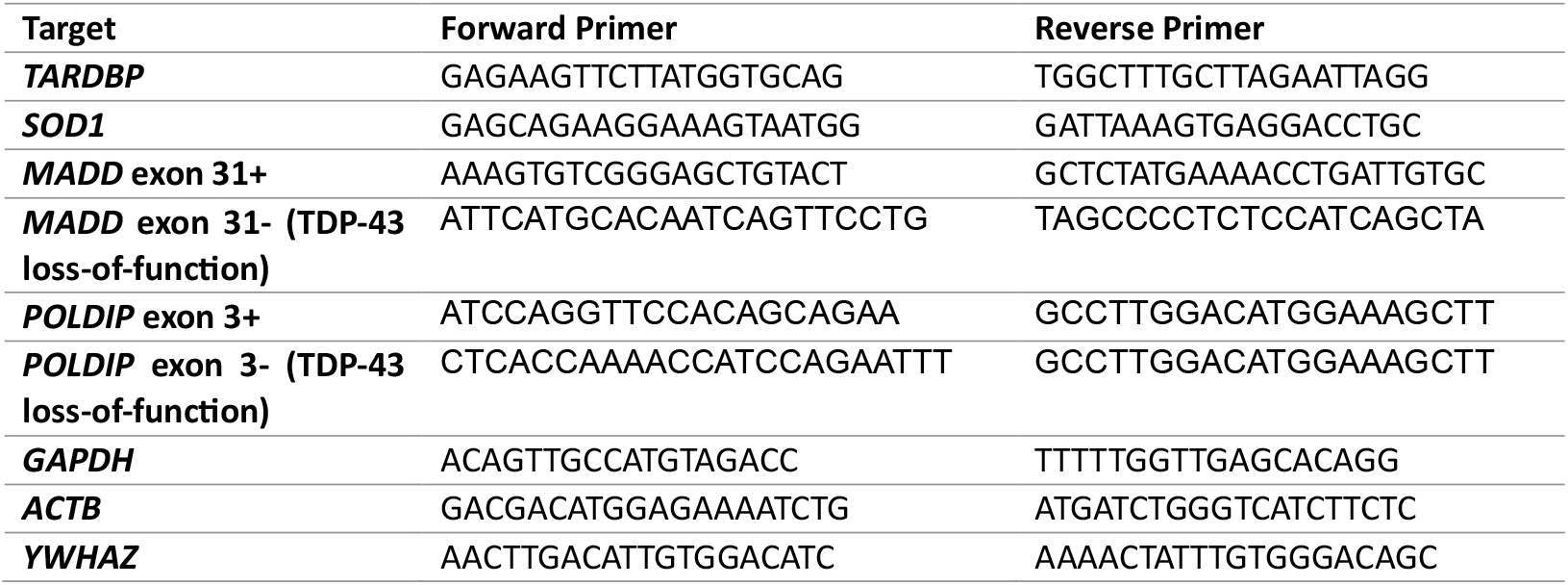
Primer sequences used for RT-qPCR.

### TDP-43 immunohistochemistry

TDP-43 immunostaining was performed using the TDP-43^APT^ (sequence: CGGUGUUGCU with a 3’ Biotin-TEG modification). The online standard operating procedure^**49**^ was followed in full, without modification. TDP-43^APT^ pathology was assessed manually by a trained histopathologist who was blinded to all demographic and clinical information. Slides were imaged using an EVOS M5000 microscope using brightfield settings at 40x magnification.

### Statistical analysis

Statistical analyses were performed using the SciPy module in python^43^. Detailed parameters are provided in the main text as appropriate, and the custom analysis code is provided as described below.

### Code & Data availability

The mass spectrometry proteomics data have been deposited to the ProteomeXchange Consortium via the PRIDE partner repository with the data set identifier PXD061429. All other pre-processed datasets and custom python scripts for all analyses are available from Zenodo (DOI 10.5281/zenodo.14960397 and 10.5281/zenodo.14965410 respectively).

## AUTHOR CONTRIBUTIONS

**Conceptualization:** DC, DK; **Funding acquisition:** DC, DK, JMG, JS; **Methodology:** DC, MB, DB; **Software:** DC; **Investigation:** DC, EL, SM, MW, FMW, JMG; **Formal analysis:** DC, EL, MW, JMG; **Visualization:** DC; **Data Curation:** DC, **Supervision:** DK, JS, JMG; **Writing - Original Draft:** DC; **Writing - Review & Editing:** all authors.

## ACKNOWLEDGMENTS

The authors are grateful to all donors, their family members and caregivers, for making this study possible. We acknowledge the MRC London Neurodegenerative Diseases Brain Bank and the Brains for Dementia Research project for facilitating the collection, storage and provisioning of donated tissues. This work was supported by the UK Dementia Research Institute, which receives its funding from DRI Ltd., funded by the UK Medical Research Council, Alzheimer’s Society, and Alzheimer’s Research. We would also like to acknowledge the support of the UK Dementia Research Institute Proteomics Platform, led by Dr Bethany Geary. D.C. was supported by the Lady Edith Wolfson Junior Non-Clinical Research Fellowship awarded by the MND Association UK (Cox 971-799). JS was supported by a Wellcome Trust Senior Research Fellowship (221824/Z/20/Z). MAW was supported by an Alzheimer’s Research UK Research Fellowship (ARUK-RF2020A-008), an award from The Sean M. Healey & AMG Center for ALS at Mass General and ALS FindingACure, and the Rosetrees Trust (CF2\100004). EL was supported by a grant from Parkinson’s UK (G-1901).

