## Supplementary Information for "Fingerprinting disease-derived protein aggregates reveals unique signature of Motor Neuron Disease"

**This file contains:**


**Supplementary Figure 1:** Schematic of tissue processing paradigm.

**Supplementary Figure 2:** TDP-43 assemblies are found in both neurologically normal and disease-derived brain extracts.

**Supplementary Figure 3:** Aggregates containing TDP-43 are found in both neurologically normal and disease brain extracts.

**Supplementary Figure 4:** TDP-43-positive assemblies differ in size and shape between neurologically normal and disease-derived brain extracts.

**Supplementary Figure 5:** Signatures of MND found in nanoscopic TDP-43 assemblies.

**Supplementary Figure 6:** Proteomic analysis of affinity-captured aggregate fraction from embedded extracts.

**Supplementary Figure 7:** Particle counts for target proteins of interest in embedded extracts.

**Supplementary Figure 8:** RT-qPCR measure of TDP-43 pathology supports SiMPull characterisation of $MND_{SOD}$ donor samples.

**Supplementary Figure 9:** Immunohistochemistry confirms evidence of TDP-43 pathology in $MND_{SOD}$ samples.

**Supplementary Figure 10:** Extracellular inflammasomes containing Apoptosis-associated Speck-like protein containing a CARD (ASC) were not altered in MND.

**Supplementary Table 1:** Qualitative summary of immunohistochemistry evidence of TDP-43 pathology.

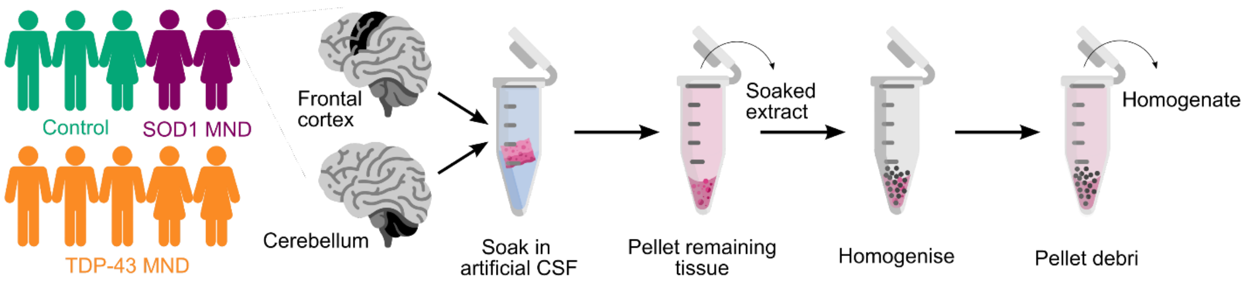


**Supplementary Figure 1: Schematic of tissue processing paradigm.** Matched frontal cortex and cerebellum samples were obtained from a cohort of 10 donors, consisting of 5 with observed TDP-43 pathology at autopsy ($MND_{TDP}$) and 5 age-matched controls lacking TDP-43 pathology, including 2 disease-control donors diagnosed with SOD1-MND ($MND_{SOD}$) and 3 neurologically normal donors (CRL). Samples were first soaked for up to 2 h in artificial CSF buffer before the remaining intact tissue was pelleted via centrifugation and the supernatant was collected. The resultant supernatant was dialysed three times in 100× the sample volume, then finally collected as the “diffusible” extract. The remaining tissue was subjected to two rounds of bead-based homogenisation after which any cellular debris were pelleted via centrifugation and the resultant supernatant collected as the “embedded” extract. These extracts were then aliquoted and stored at -80 °C until use.

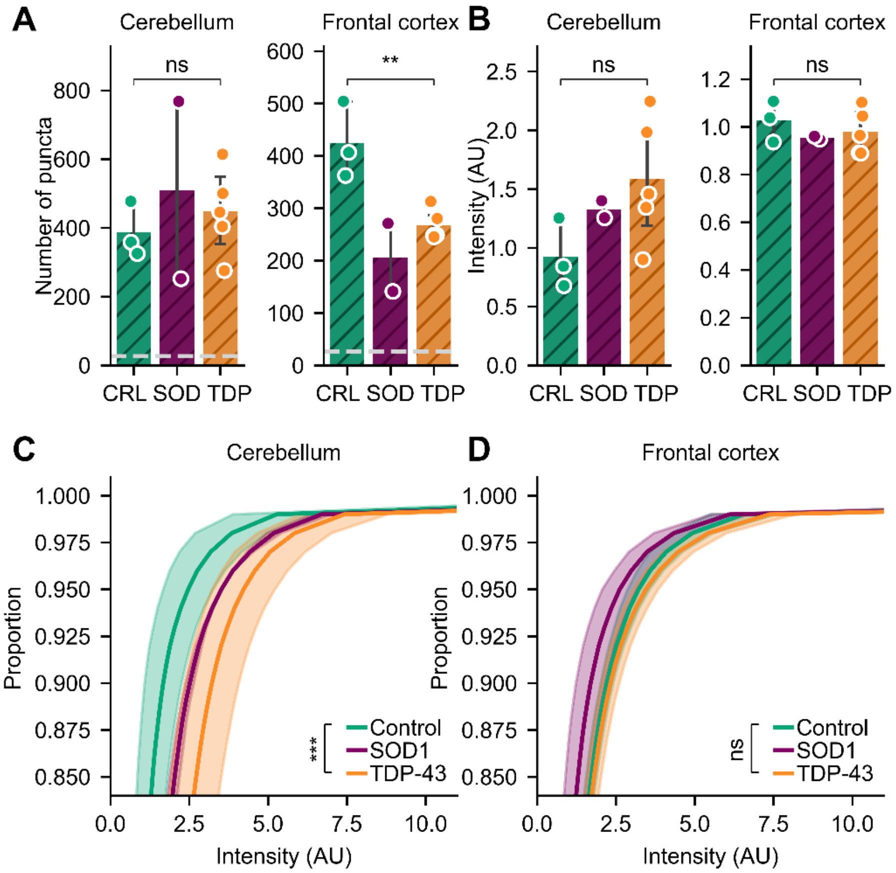


**Supplementary Figure 2: TDP-43 assemblies are found in both neurologically normal and disease-derived brain extracts. (A)** Number and **(B-D)** brightness of TDP-43 positive assemblies captured with flTDP-SiMPull from embedded extracts. Shown in *A*, *B* are the mean ± S.D. of donors in each category overlayed with the raw mean datapoints from three technical replicates of each donor, compared using Welch's t-test, ** $p < 0.01$, ns $p > 0.05$. Dashed line corresponds to an isotype control exposed to a reference sample of the same extract type. Shown in *C*, *D* are the mean interpolated cumulative distributions for each cohort ± S.D. of individual donors, compared via Kolmogorov–Smirnov test with Bonferroni correction, *** $p < 0.001$, ns $p > 0.05$.

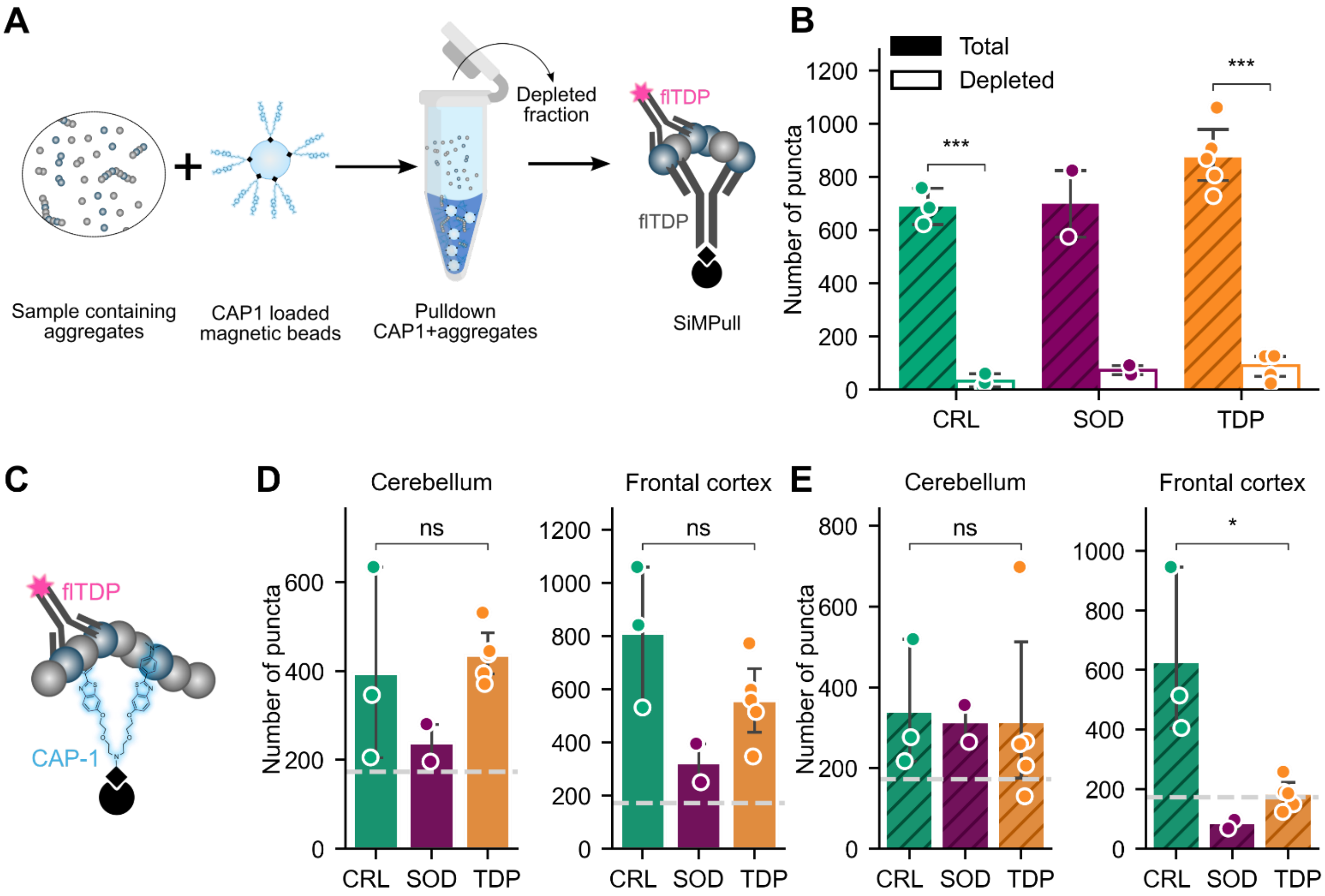


**Supplementary Figure 3: Aggregates containing TDP-43 are found in both neurologically normal and disease brain extracts. (A)** Schematic of CAP1-bead based aggregate affinity pulldown applied to embedded extracts followed by quantification via TDP-43-SiMPull. **(B)** Number of TDP-43 assemblies before (total) and after (depleted) CAP1 affinity capture of aggregates from the sample. Shown is mean ± S.D. of each cohort, overlayed with the raw mean datapoints from each donor, compared via two-way ANOVA ($p < 0.0001$) with post-hoc t-test with Holm-Bonferroni correction, *** $p < 0.001$. **(C)** Schematic of the CAP1-SiMPull setup applied to detect TDP-43 positive particles in *D, E*. **(D-E)** Number of TDP-43 positive particles in *(D)* diffusible and *(E)* embedded extracts captured with CAP1-SiMPull. Shown are the mean ± S.D. of each cohort, overlayed with the raw mean datapoints from three technical replicates of each donor. Dashed line corresponds to an isotype control exposed to a reference sample of the same extract type. Comparisons derived from Welch's t-test, * $p < 0.05$.

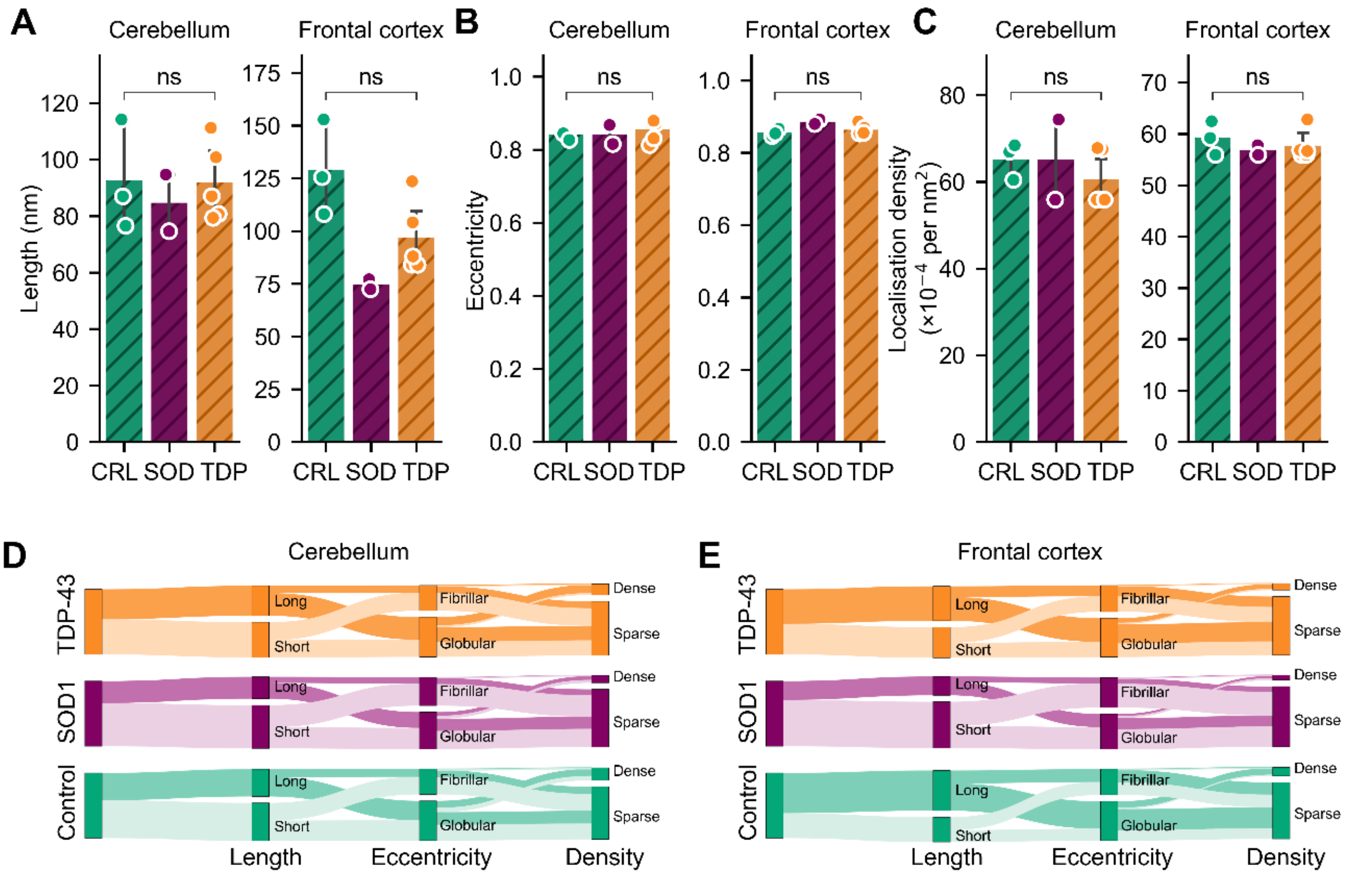


**Supplementary Figure 4: TDP-43-positive assemblies differ in size and shape between neurologically normal and disease-derived brain extracts. (A-C)** Average **(A)** length, **(B)** eccentricity, and **(C)** localisation density of TDP-43 positive assemblies captured with CAP1-SiMPull from embedded extracts. **(D-E)** Sankey visualisation of the proportion of assemblies that fall into each category as defined by cumulative thresholds for length (long > 100 nm), eccentricity (fibrillar > 0.9) and localisation density (dense > 0.001 localisations per nm$^2$). Shown in *A-C* are the mean ± S.D. of donors in each category overlayed with the raw mean datapoints from three technical replicates of each donor, compared using Welch's t-test. Shown in *D*, *E* are the compiled assembly populations for each cohort.

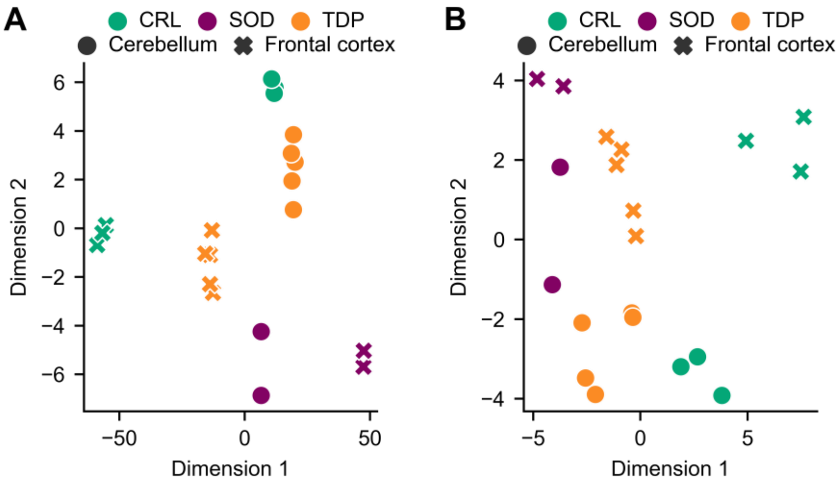


75 **Supplementary Figure 5: Signatures of MND found in nanoscopic TDP-43 assemblies.** Two-dimensional projection of linear discriminant analysis performed on (**A**) particle properties derived from embedded extracts or (**B**) combined particle properties from diffusible and embedded extracts characterised via SiMPull.

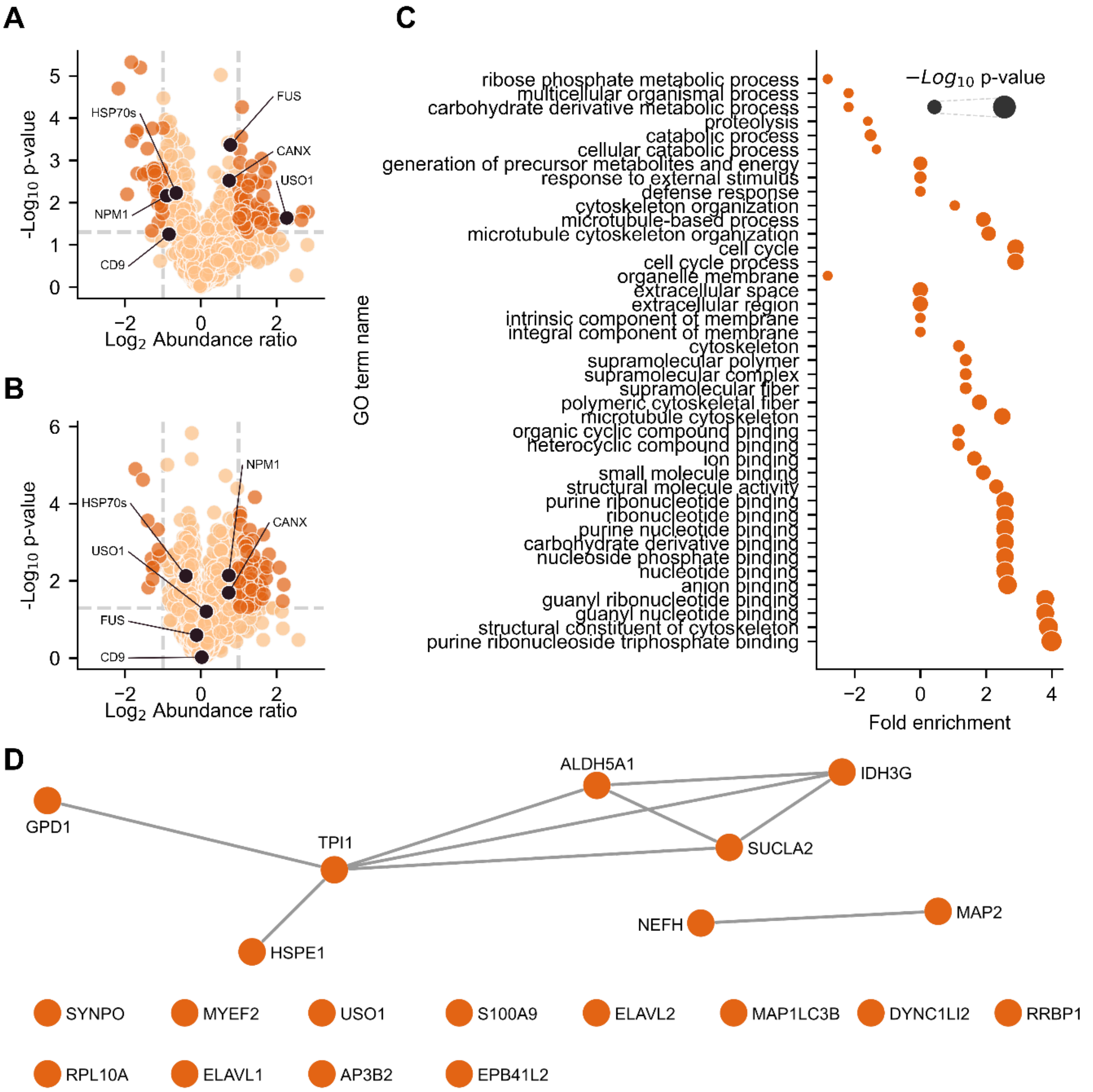


**Supplementary Figure 6: Proteomic analysis of affinity-captured aggregate fraction from embedded extracts. (A-B)** Volcano plot depicting abundance ratio of proteins identified in **(A)** the CAP1 aggregate fraction and **(B)** the total proteome from frontal cortex embedded extracts of MND$_{TDP}$ vs neurologically normal control cohorts. Thresholds depict $|log_2FC| > 1$ and $-log_{10}(p-value) > 1.3$ outside which proteins are considered significantly altered (dark orange). Proteins of interest (black) highlighted. **(C)** Panther GOSLIM gene ontology enrichment and **(D)** STRINGdb protein-protein interaction network for proteins enriched in the CAP1 aggregate fraction from MND$_{TDP}$ embedded extracts. Nodes represent individual proteins connected.

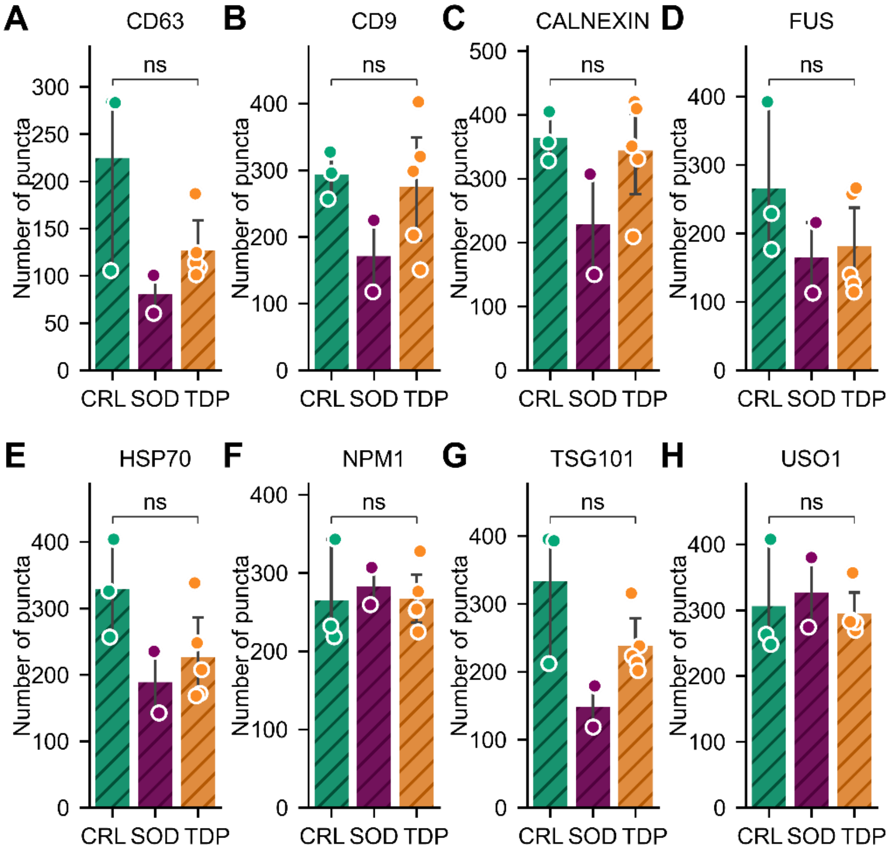


**Supplementary Figure 7: Particle counts for target proteins of interest in embedded extracts.** Number of aggregates captured with CAP1-SiMPull positive for **(A)** CD63, **(B)** CD9, **(C)** Calnexin, **(D)** FUS, **(E)** HSP70, **(F)** NPM1, **(G)** TSG101 and **(H)** USO1. Shown are the mean ± S.D. of donors in each category overlayed with the raw mean datapoints from three technical replicates of each donor compared using Welch's t-test, ns $p > 0.05$.

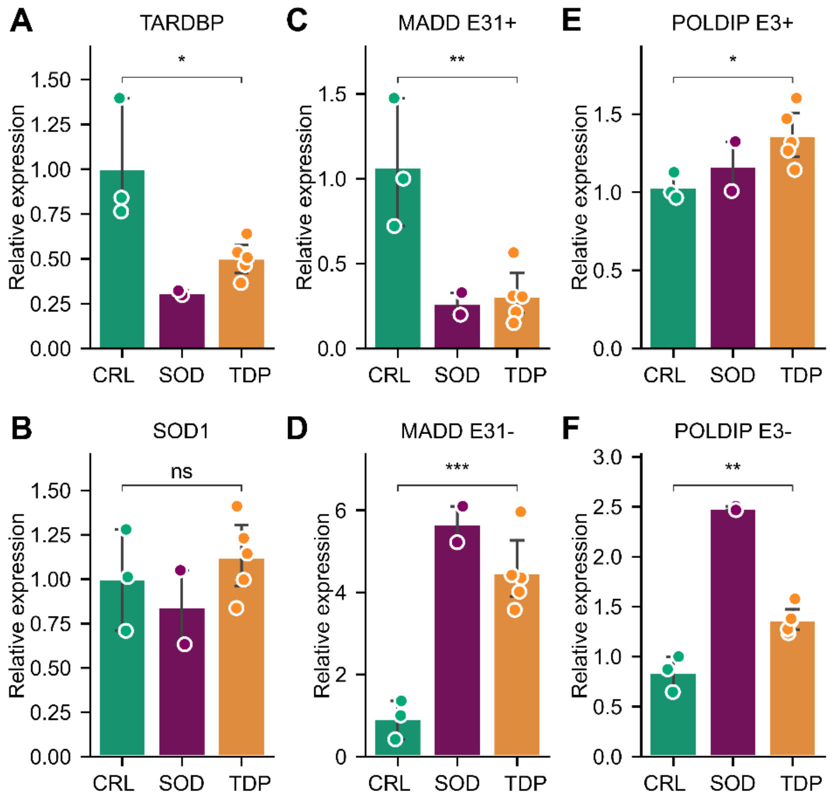


**Supplementary Figure 8: RT-qPCR measure of TDP-43 pathology supports SiMPull characterisation of MND$_{SOD}$ donor samples. (A-F)** Relative expression of six target genes in frontal cortex samples as measured by quantitative PCR (qPCR). Gene counts were normalised to reference genes *GAPDH*, *ACTB* and *YWHAZ.* Expression of canonical **(A)** TDP-43 and **(B)** SOD1 is shown, alongside two TDP-43 splicing targets **(C-D)** *MADD* and **(E-F)** *POLDIP* for which inclusion (+) of specific exons (E31 and E3 respectively) yields the canonical transcript, while exclusion (-) is associated with loss of TDP-43 function. Shown are the mean ± S.D. of donors in each category overlayed with the raw mean datapoints from three technical replicates of each donor compared using Welch's t-test, ns $p > 0.05$ * $p < 0.05$ ** $p < 0.01$ *** $p < 0.001$.

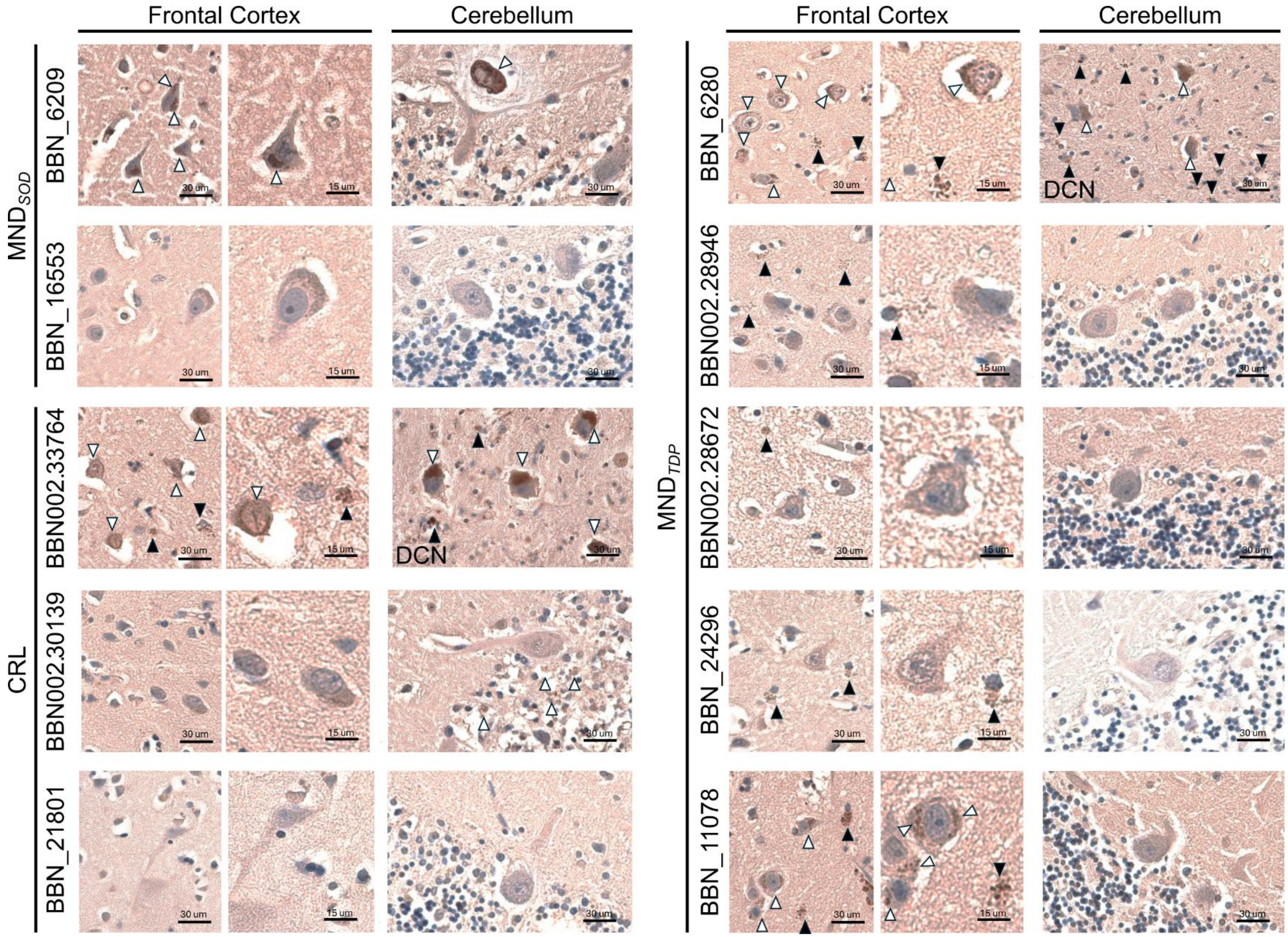


**Supplementary Figure 9: Immunohistochemistry confirms evidence of TDP-43 pathology in MND$_{SOD}$ samples.** Immunohistochemical staining performed using TDP-43 aptamer, used to detect, with high sensitivity, TDP-43 aggregates in both neurons and glia. Shown are exemplar images for each donor stained with TDP-43$_{APT}$[50,51] from frontal cortex and cerebellar sections. White arrows indicate neuronal TDP-43 pathology, black arrows indicate glial TDP-43 pathology, DCN indicates neurons within the deep cerebellar nuclei, and individual scale bars are displayed for each image. See Supplementary Table 1 for summarised qualitative findings of TDP-43 pathology.

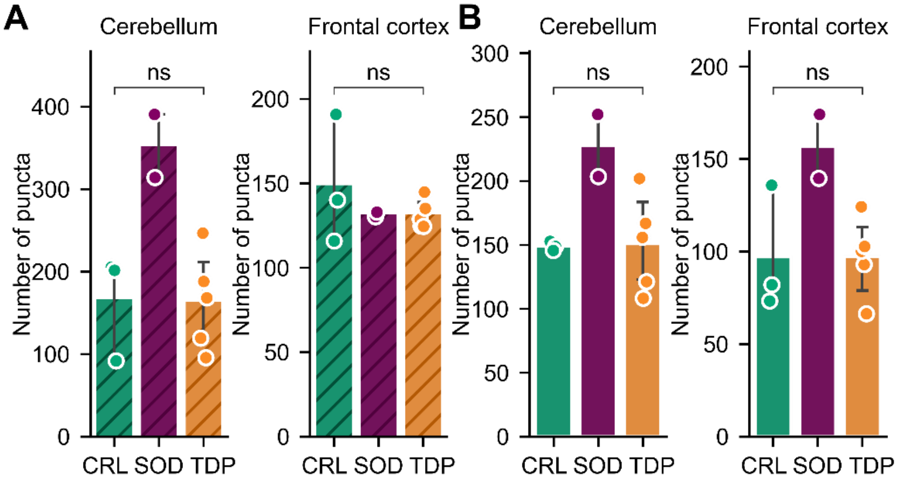


**Supplementary Figure 10: Extracellular inflammasomes containing Apoptosis-associated Speck-like protein containing a CARD (ASC) were not altered in MND. (A-B)** Number of ASC positive assemblies[39] captured from **(A)** embedded and **(B)** diffusible extracts. Shown are the mean ± S.D. of donors in each category overlayed with the raw mean datapoints from three technical replicates of each donor compared using Welch's t-test, ns $p > 0.05$.

**Supplementary Table 1: Qualitative summary of immunohistochemical staining with the TDP-43 aggregate aptamer by qualified pathologist.** See Supplementary Figure 9 for exemplar images highlighting TDP-43 pathology.

| MRC BNN-ID | Category | TDP-43 pathology (TDP-43$_{APT}$) | |
|---|---|---|---|
| | | Frontal Cortex | Cerebellum |
| BNN_6209 | SOD1-MND | Abundant cytoplasmic TDP-43 pathology in neurons. No apparent glial pathology. | Heterotopic Purkinje cell containing abundant cytoplasmic TDP-43 pathology. |
| BBN_16553 | SOD1-MND | No pathology. | No pathology. |
| BNN002.28672 | TDP43-MND | Occasional glial TDP-43 pathology, no neuronal TDP-43 pathology. | No pathology. |
| BNN_24296 | TDP43-MND | Abundant glial TDP-43 pathology, no neuronal TDP-43 pathology. | No pathology. |
| BNN_11078 | TDP43-MND | Abundant neuronal and glial TDP-43 pathology. | No pathology. |
| BNN_6280 | TDP43-MND | Abundant neuronal and glial TDP-43 pathology. | No TDP-43 pathology in grey matter but extensive TDP-43 pathology in deeper structures of the cerebellum (i.e. in cerebellar nuclei). |
| BNN002.28946 | TDP43-MND | Occasional glial TDP-43 pathology. | No pathology. |
| BNN002.33764 | Control | Abundant neuronal and glial TDP-43 pathology including nuclear and cytoplasmic pathology. | No TDP-43 pathology in grey matter but extensive TDP-43 pathology in deeper structures of the cerebellum (i.e. in cerebellar nuclei). |
| BNN002.30139 | Control | No pathology. | Occasional TDP-43 pathology noted in granular cell layer. |
| BNN_21801 | Control | No pathology. | No pathology. |
